# An *in vitro* tumorigenesis model based on live cell-generated oxygen and nutrient gradients

**DOI:** 10.1101/2020.08.24.264580

**Authors:** Anne C. Gilmore, Sarah J. Flaherty, Veena Somasundaram, David A. Scheiblin, Stephen J. Lockett, David A. Wink, William F. Heinz

## Abstract

The tumor microenvironment (TME) is multi-cellular, spatially heterogenous, and contains cell-generated gradients of soluble molecules. Current cell-based model systems lack this complexity or are difficult to interrogate microscopically. We present a 2D live-cell chamber that approximates the TME and demonstrate that breast cancer cells and macrophages generate hypoxic and nutrient gradients, self-organize, and have spatially varying phenotypes along the gradients, leading to new insights into tumorigenesis.

---

Concentration gradients of soluble molecules in tissue are established by their release from and consumption by cells in combination with extracellular diffusion. These gradients influence the spatial organization and phenotypes of cells in solid tissues, including tumors^1^.

Current experimental tissue models do not capture the complex spatial organization of cells and molecules, and they can be difficult to interrogate microscopically. Standard 2D cell culture does not replicate long-range (> 100 μm) gradients of extracellular molecules within the TME. Microfluidic systems can impose diffusive gradients on 2D cell cultures and are designed for microscopic interrogation^2, 3^, but they control only a limited number of molecules of interest, the spatial organization of cells is not the same as tissue, and the molecular gradients are not naturally cell driven. Although organoids and spheroids reflect to some extent the molecular gradients that arise in actual tissues, they vary in structure^4^ and are challenging to examine and quantify at high spatial resolution with long term live-cell microscopy^5^.

Hypoxic gradients in 2D cell culture were observed in 2018 in restricted exchange environment chambers (REECs)^6^. Cells in this chamber, supported on a standard #1.5 glass coverslip (0.17 mm thick), grow in a small (< 20 μL) lower compartment separated from a larger upper compartment (^~^ 1 mL) by a coverslip with a small central hole (^~^0.7 mm diameter) through which O_2_ and soluble molecules (e.g., nutrients, cytokines, and cellular waste products) diffuse between the compartments (Fig. 1a). Thus, cells directly beneath the hole are exposed to O_2_ and nutrients at the concentration of the upper compartment, while cells distal to the hole exist in a cell-generated hypoxic environment.

**Figure 1.**
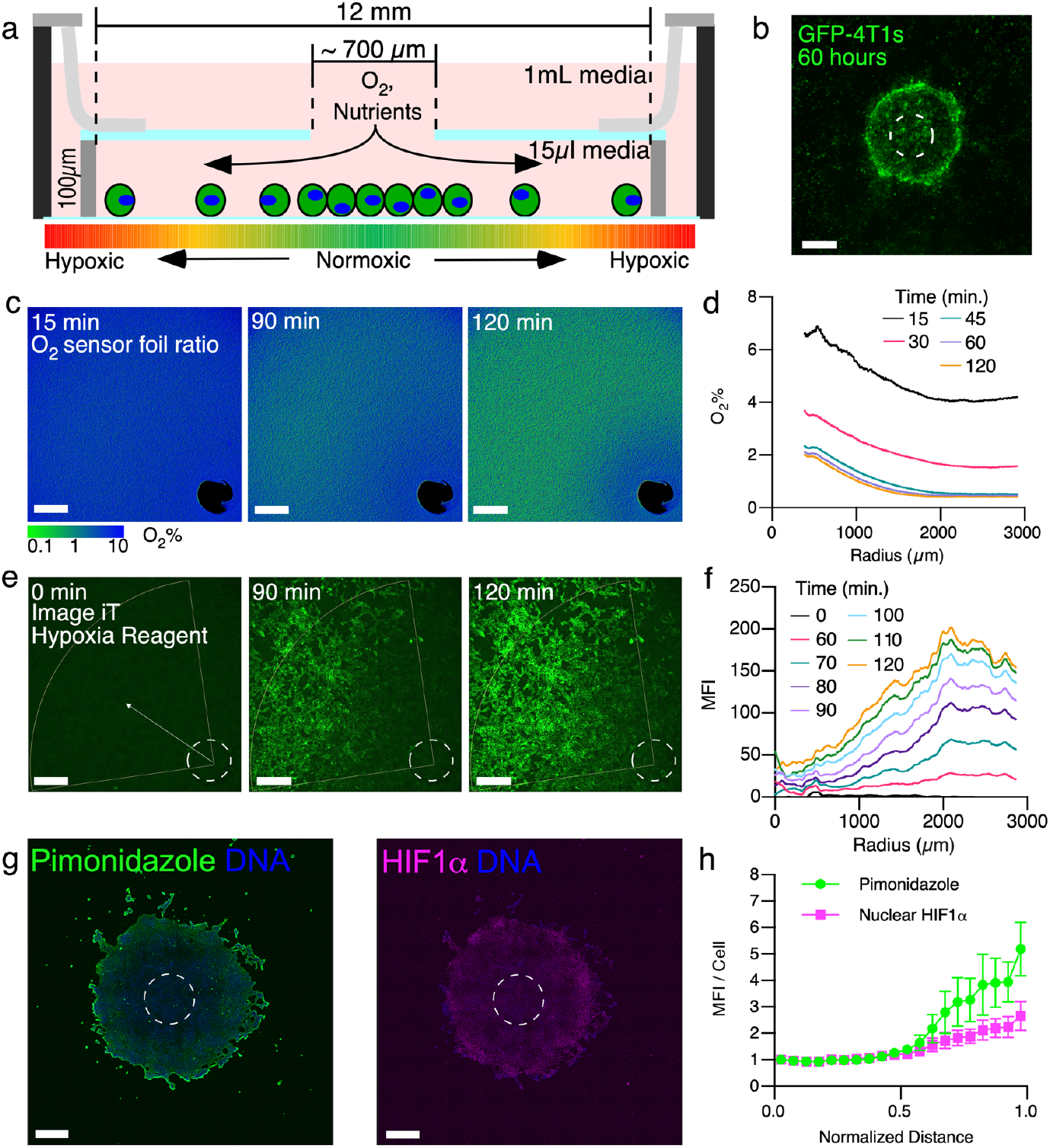
In the 4T1-REEC model of the TME, cell-generated [O_2_] gradients correlate with intracellular hypoxia. a. Schematic of restrictive exchange environment chamber (REEC). b. GFP-expressing 4T1 cells migrate up the oxygen gradient to form a disk centered around the hole in the REEC (scale bar = 500 μm). c. Cell-generated extracellular [O_2_] gradients in a 4T1-REEC. Ratiometric images of an O_2_ sensor foil over two hours (scale bar = 500 μm). d. Quantification of extracellular [O_2_] at various time points for the 4T1-REEC in c. e. Intracellular hypoxic gradients. Widefield images of Image IT Hypoxia Green Reagent (IHGR), which fluoresces in response to intracellular hypoxia, in a 4T1-REEC over two hours (scale bar = 500 μm). f. MFI of IGHR versus distance from opening at various time points. g. Widefield image of pimonidazole (left) and HIF1α (right) immunofluorescence staining of 7-day old 4T1 disk (scale bar = 500 μm). h. Quantification of pimonidazole and nuclear HIF1α immunofluorescence in 4T1 disks as a function of distance from opening to the edge of the disk (N=7 disks). MFI per cell is normalized to the first point. Dashed lines indicate the opening of the hole. MFI = mean fluorescence intensity.

Here we report the application of the chamber to investigate the TME. Specifically, we examine the role of O_2_ and nutrient gradients on tumor and immune cell phenotype, using 4T1 mouse mammary tumor cells and ANA-1 mouse macrophages as exemplars. The 4T1 model shares many features with triple negative human breast cancer^7^, and macrophages are the most abundant non-cancer cells in the TME and play an immunosuppressive role in tumorigenesis^8^. These cells types were cultured separately and in co-culture in the chamber, and the spatiotemporal dynamics of O_2_ gradient formation, nutrient uptake, cell migration and cell survival were quantified.

## Results

GFP-tagged 4T1 cells, initially uniformly distributed (^~^75% confluence) in the lower compartment, began migrating towards the hole within 36 h and formed a stable disk with a diameter of ^~^1 mm within ^~^60 h (Fig. 1b, Supplemental Video 1). Beyond the disk, cells detached from the bottom of the well and died (Supplementary Fig. 1), analogous to necrotic zones in solid tumors. The resulting 4T1-disk remained alive and stable for at least three weeks with periodic changes to media in the upper compartment. We attribute the directed motion of cells to O_2_ gradients within the chamber, which do not arise in standard cell culture and cannot be readily observed in intact tumors or 3D cell culture models. 4T1 proliferation recovers with disk expansion after the chamber is removed (Supplemental Fig. 2) and normoxia is reestablished. Interestingly, this experiment showed that 4T1 cells could respond to the gradient of O_2_ by converting to a migratory phenotype in order to coalesce near the opening in the chamber. This behavior is characteristic of tumor cells *in vivo* and was shown here to be possible without the presence of other cell types, extracellular matrix, or 3D environment.

After demonstrating that REECs reproduce tumor behavior, we quantitatively characterized the hypoxia and nutrient gradients, and how these gradients affected 4T1 phenotype and metabolism. Dissolved extracellular O_2_ concentration ([O_2_]) measurements in the REECs showed that stable radial negative gradients formed rapidly (< 2 h) after chamber placement onto uniformly distributed 4T1 cells in 2D cell culture (Fig. 1c, d; Supplemental Fig. 3), which agreed with mathematical modeling (Supplemental Fig. 4). Fluorescence labeling of the live 4T1 cells with the image-iT green hypoxia reagent (IGHR) revealed that positive gradients of intracellular hypoxia formed in a similar timescale, and the hypoxic front (the distance at which the IGHR signal is 90% of its maximum) was within a millimeter of the edge of the opening after 2 h (Fig. 1e, f).

Hypoxia, through hypoxia inducible factor (HIF1α) stabilization, can induce pro-tumor phenotypes (Supplemental Fig. 5), including a collective-to-amoeboid transition, increased in vimentin expression, and migration of 4T1 cells^9^. Cells in the periphery of the disks exhibit a temporary increase in vimentin expression relative to E-cadherin expression, between 12 and 36 hours after initiation, suggesting a transient phenotypic shift to a mesenchymal state (Supplemental Fig. 6)^10^. In stable disks, increased cellular hypoxia distal to the hole was measured by pimonidazole reduction and HIF1α localization to the nuclei (Fig. 1g, h).

During initial O_2_ gradient formation (< 2 h) glucose consumption exhibited a radial gradient with maximum consumption at the opening at early time points (Fig. 2a, b;). However, no gradient of mitochondrial membrane potential (ΔΨm) was observed via tetramethylrhodamine-ethyl-ester (TMRE) fluorescence 2 hours after chamber placement, suggesting that oxidative phosphorylation continues for some time after hypoxia is established. In fully formed disks (> 72 h), ΔΨm decreased while glucose consumption increased with radial distance (Fig. 2c, d, Supplemental Fig. 7), consistent with a metabolic shift from oxidative phosphorylation to glycolysis in the hypoxic region. Mathematical modeling predicted no significant gradient of glucose in the media away from the standard serum glucose concentration (22.5 mM), which is far in excess of the metabolic needs of the cells (Supplemental Fig. 4). Thus, we conclude the [O_2_] gradient alone drove the metabolic shift. These results demonstrate that REECs capture key features of hypoxia and metabolism in 4T1 tumorigenesis.

**Figure 2.**
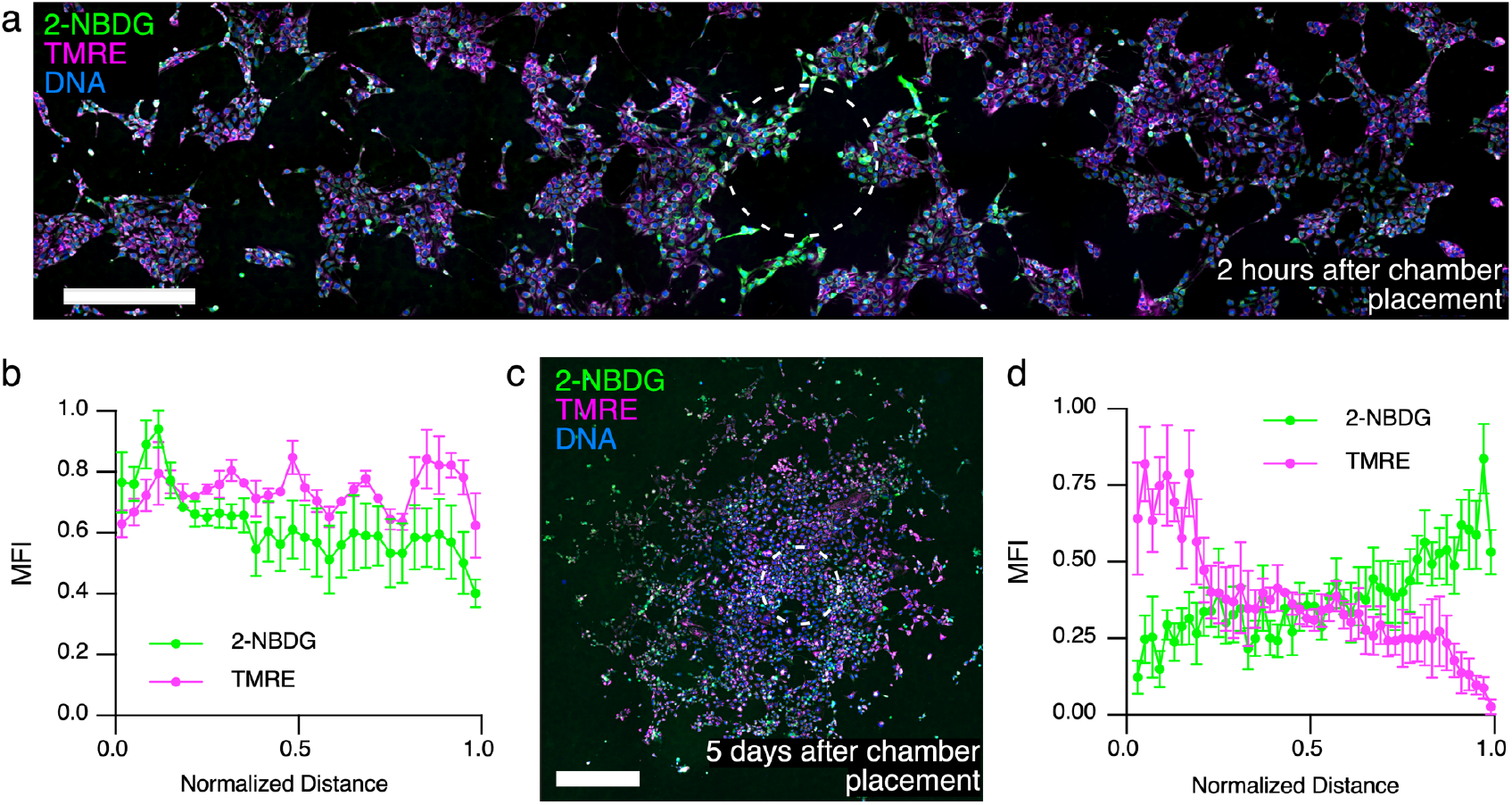
Nutrient uptake and metabolic activity vary with [O_2_] concentration gradient between 2 hours and 5 days in the 4T1-REEC model of the TME. a. Widefield of image of 2-NBDG uptake and TMRE fluorescence in a 4T1-REEC 2 hours after chamber placement (scale bar = 500 μm). b. Quantification of 2-NBDG and TMRE fluorescence in 4T1-REECs 2 hours after chamber placement (N = 3). Correlation coefficient of 2-NBDG, TMRE = 0.1748. c. Widefield image of 2-NBDG and TMRE fluorescence in a 4T1-REEC 5 days after chamber placement (scale bar = 500 μm). d. Quantification of 2-NBDG (N = 6 disks) and TMRE (N = 3 disks) fluorescence in 4T1-REECs 5 days after chamber placement. Correlation coefficient of 2-NBDG, TMRE = −0.5089. Dashed lines indicate the opening of the hole. MFI = mean fluorescence intensity.

We utilized REECs to further understand the immunosuppressive role of the inflammatory proteins inducible nitric oxide synthase (Nos2) and cyclooxygenase-2 (Cox2) in the tumorigenic microenvironment. Clinically, high expression of both these proteins in ER-human breast cancer is an indicator of very poor prognosis^11^; both are potential targets for therapy using FDA-approved anti-inflammatory drugs in combination with standard treatments. Nos2 produces nitric oxide (NO), a key regulator of cancer processes^12^, from L-arginine and is upregulated by stabilized HIF1α in response to hypoxia and nutrient deprivation^13^ (Supplemental Fig. 5). Therefore, we postulated that in the hypoxic regions of fully formed disks, the Nos2 expression and NO flux would be high relative to Cox2-expression. We observed Nos2 levels remain high relative to Cox2 in the hypoxic regions of disks, spheroids, and tissue (Fig. 3a, b, Supplemental 8). We also saw a large increase in NO flux in the hypoxic regions of 4T1-disks, confirming our hypothesis (Figure 4a-d, control disks; Supplemental 8).

**Figure 3.**
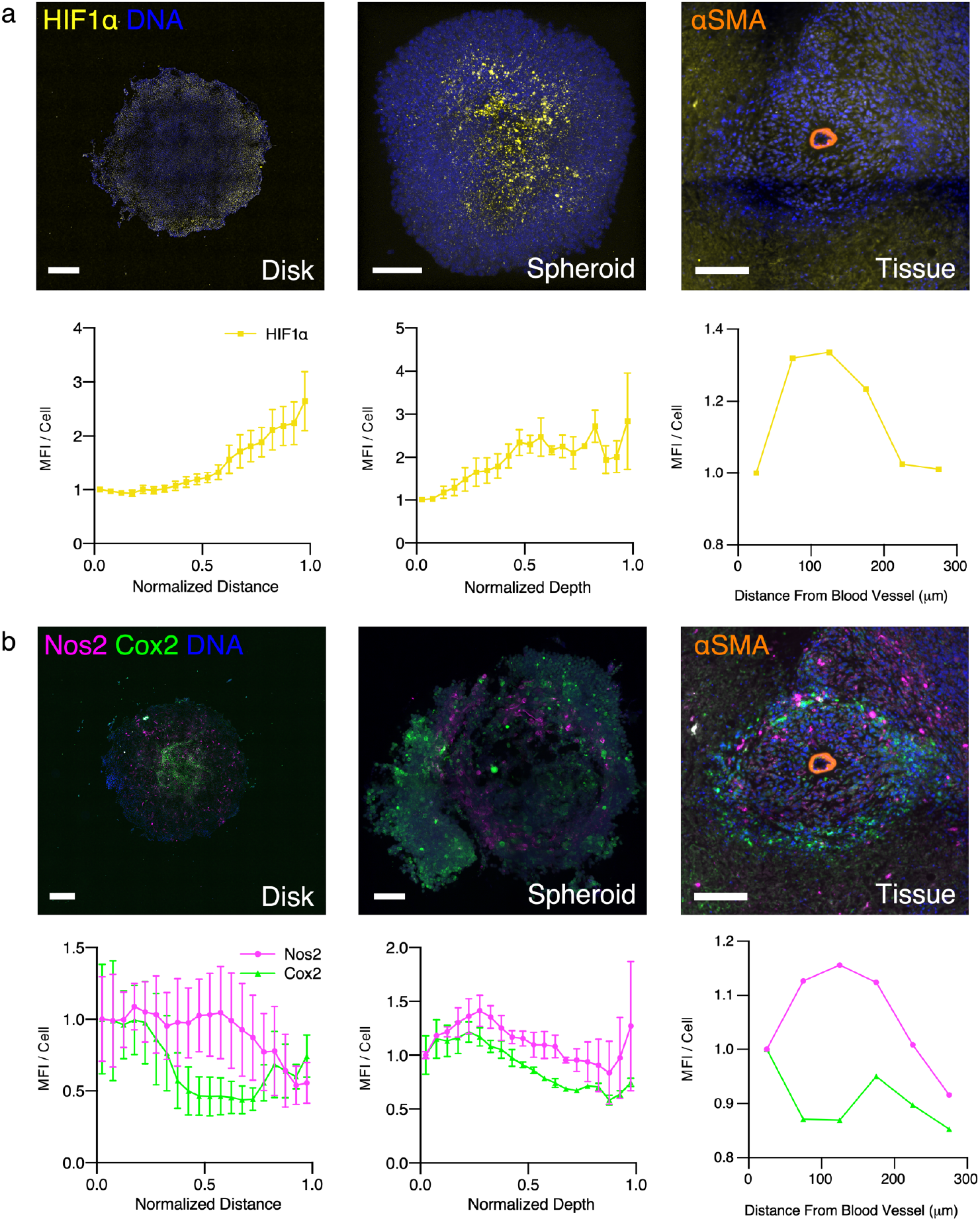
Spatial distribution of cellular phenotypes along hypoxic gradients in 4T1-REECs, 4T1 spheroids, and 4T1 tumor tissue are similar. a. Distributions of HIF1α fluorescence in 4T1 disks (N=7), spheroids (N=4), and tissue (N=1). b. Distributions of Nos2 and Cox2 fluorescence in 4T1 disks (N=3), spheroids (N=2), and tissue (N=1). The tissue images capture a cross section of a blood vessel. Immunofluorescent staining of α-smooth muscle actin (αSMA) labels the vascular smooth muscle cells. “Normalized Distance” refers to the relative distance from the center to the edge of a disk. “Normalized Depth” refers to relative distance from the surface to the center of a spheroid. “Distance from Blood Vessel” is measured from the center of the blood vessel. MFI = mean fluorescence intensity normalized to the first point. Disk scale bar = 500 μm. Spheroid and tissue scale bars = 100 μm.

**Figure 4.**
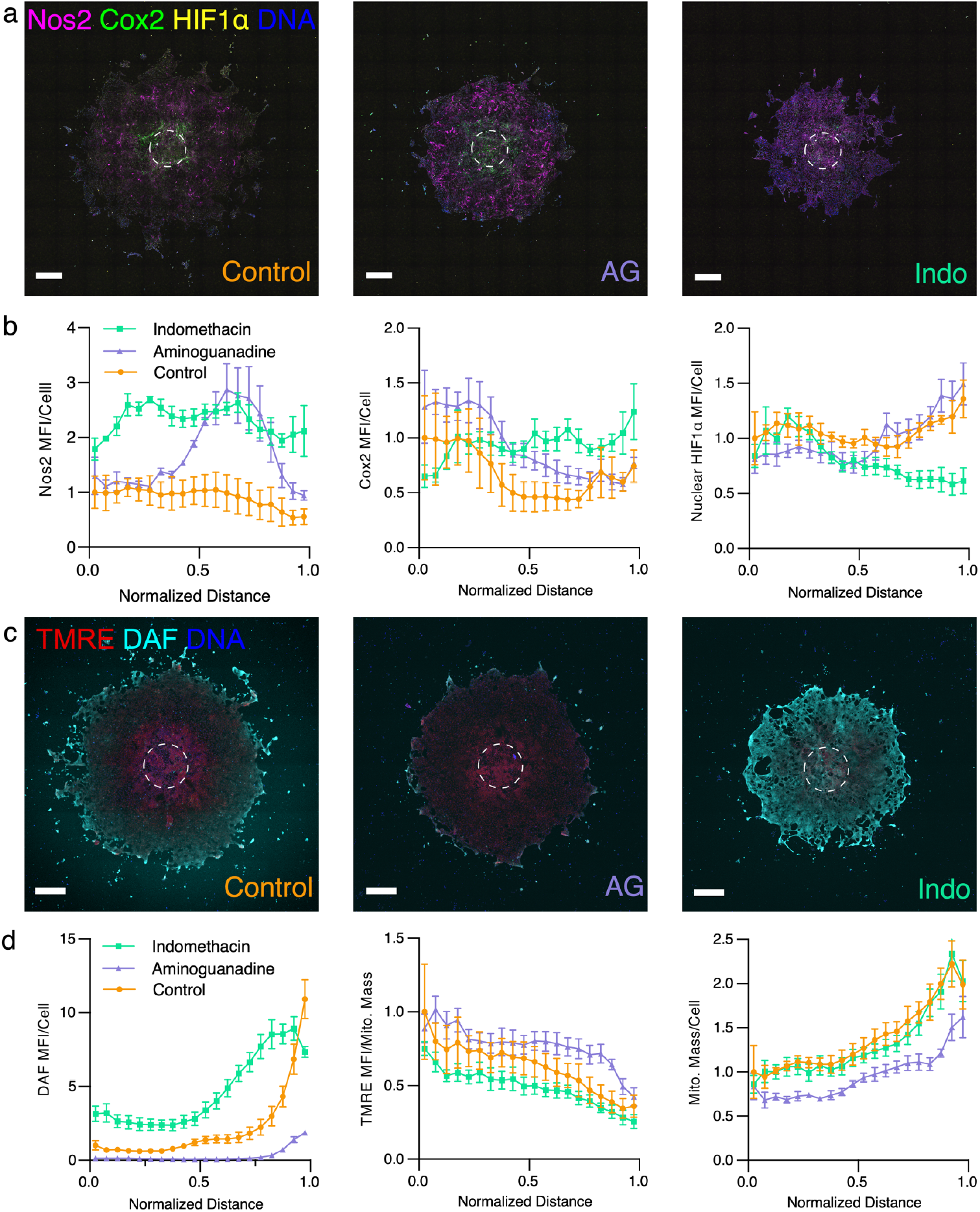
Effects of Nos2 inhibition by aminoguanidine (AG, 1 mM) or Cox2 inhibition by indomethacin (Indo, 100 μM) on 4T1 phenotype along hypoxic gradients. a. Spatial distribution of Nos2, Cox2, and HIF1α imaged by widefield immunofluorescence microscopy of 7-day old untreated 4T1 disks (control), AG-treated disks, and Indotreated disks. b. Nos2, Cox2, and HIF1α distributions in 7-day old untreated 4T1 disks (control, N = 3), AG-treated disks (N = 4 for Nos2, Cox2, & HIF1α), and Indo-treated disks (N = 4). c. Spatial distribution of DAF and TMRE fluorescence measured by confocal microscopy of 7-day old untreated 4T1 disks (control), AG-treated disks, and Indo-treated disks. d. DAF, TMRE, and ATP5A distributions in 7-day old untreated 4T1 disks (control, N = 4), AG-treated disks (N = 4), and Indo-treated disks (N = 4). “Normalized Distance” refers to the relative distance from the center to the edge of a disk. MFI = mean fluorescence intensity normalized to the first point of the untreated control. Scale bars = 500 μm.

We observed in disks, spheroids, and tumor tissue that different 4T1 cells expressed high levels of Nos2 or Cox2 and that high Nos2-expressing cells tended to be clustered together (Fig. 3b, Supplemental Fig. 9) under hypoxia-induced stress conditions. The fact that Nos2 and Cox2 are highly expressed in different cells indicates that these proteins likely drive the expression and activity of each other via an intercellular feed-forward mechanism^14^ mediated by the release of NO and PGE2 from the cells^11^. Nos2-expressing cell clumps likely lead to significantly higher local concentrations of NO than could arise when Nos2-expressing cells are scattered and isolated^15^ and thus augment this paracrine mechanism.

Inhibitors of Nos2 and Cox2 interrupt the feed-forward loop and reduce tumor growth rate^11, 13^. We therefore treated 4T1 disks in REECs with Nos2 or Cox2 inhibitors for 7 days to examine the effect of the inhibitors on the distribution of protein expression, NO release, ΔΨm, and mitochondrial mass within the disks (Fig. 4a-d).

Treatment with the Nos2 inhibitor aminoguanidine (AG, 1 mM) resulted in a large compensatory increase in Nos2 expression in the more hypoxic regions of the disk versus the untreated control, minimal NO levels, as well as increased Cox2 expression in the more normoxic regions of the disk. Treatment with the Cox inhibitor indomethacin (Indo, 100 μM) likewise resulted in a compensatory increase in Cox2 expression in hypoxic regions of the disk and increased Nos2 expression and NO levels across the disk relative to control. Interestingly, nuclear HIF1α levels were lower in the hypoxic regions of Indo-treated disks relative to the controls (Fig. 4a, b). Given that NO flux increases in the hypoxic region with indomethacin treatment (Fig. 4c, d), this is most likely due to the known inhibitory effect of NO on HIF1α in hypoxic conditions^16^.

ΔΨm was higher in more normoxic regions compared to hypoxic regions for both treatments and controls, as expected (Fig. 4c, d). Relative to controls, ΔΨm was lower in Indo-treated disks, indicating decreased mitochondrial activity in response to the higher levels of NO across the disk, as expected. ΔΨm was higher in AG treated disks, particularly in the hypoxic regions, indicating a failure to switch to anaerobic metabolism in absence of functional Nos2. Additionally, mitochondrial mass was lower in the AG treated disks (Fig. 4d), likely due to the lack of NO, which plays an important role in mitochondrial biogenesis^17^. Indo-treated 4T1 disks contained fewer cells than control disks, mirroring effects of the anti-inflammatories in tumors (Supplementary Fig. 10)^12,13^. These results show that anti-inflammatory compounds modulate cellular phenotypes along the hypoxic gradient, which demonstrates the utility of the 4T1-REEC system for understanding treatment mechanisms in conjunction with hypoxia in the TME.

Macrophages localize to areas of hypoxia and necrosis in the TME where they play an immunosuppressive role^18^. Therefore, we cultured ANA-1 macrophages in the REEC either alone or with 4T1 cells. The macrophages, unlike 4T1 cells did not migrate towards the opening or form disks, though they did generate a hypoxic gradient as observed via IGHR staining (Supplemental Fig. 11). Macrophages treated with the pro-inflammatory cytokines IFNγ and LPS express higher levels of Nos2 and thus produce more NO, which converts them to a glycolytic pathway and decreases their mitochondrial metabolism and O_2_ consumption^15^. This effect was observed in REECs as the hypoxic front of treated macrophages was further from the hole than that of untreated cells (Supplemental Fig. 11). When macrophages were co-cultured with 4T1s or injected through the hole onto stable 4T1 disks, the macrophages populated the hypoxic and 4T1-necrotic regions (Supplemental Fig. 11). These results are consistent with macrophage behavior in spheroids and *in vivo* in which IFNγ+LPS stimulated macrophages infiltrate the hypoxic core of spheroids (Supplemental Fig. 12) and hypoxic regions of tumors^19^. Interestingly, the area of stable 4T1 disks decreased following the injection of unstimulated macrophages. Injection of stimulated macrophages or fresh media resulted in increases in 4T1 disk area (Supplemental Figure 11). This suggests that the stimulated macrophage-generated increase in extracellular NO in the chamber drives glycolysis and reduces the O_2_ consumption of the 4T1 and ANA1 cells enough to push back the hypoxic front, thereby promoting survival and proliferation of the tumor cells.

In conclusion, we demonstrated that the 4T1/ANA1-REEC *in vitro* model captures key features of the tumorigenic microenvironment. It recapitulates the cell-generated oxygen gradients that exist in solid tumors and via live-cell microscopy can reveal cell dynamics and phenotypes that cannot be readily determined from actual tumors. Hence, the REEC system is a powerful tool to investigate mechanisms of tumorigenesis, immunotherapy, and anti-inflammatory treatments in live cells in a tumor-like environment.

## Methods

### REEC design and assembly

The REECs design was modified to improve functionality and fit 12-well glass-bottom plates (Figure 1A). For each REEC, a through hole (^~^700 μm diameter) was manually machined into a circular cover glass (18 mm diameter) using a high-speed air drill and tapered carbide bit (SCM Systems, Inc, Menomonee Falls, WI). A stainless-steel O-ring (0.100 mm thick, 12 mm ID, 18 mm OD) was epoxied to one side of the cover glass using UV-curable epoxy (Norland Products, Inc.). Laser-machined mylar clamps were epoxied to the other side of the cover glass using the same epoxy. Between each step, the epoxy was cured for 5 minutes in a UV-Ozone cleaner (Model 342, Jelight). Chambers were UV-sterilized immediately prior to use in cell culture (Supplemental Protocol).

The stainless-steel O-ring has as smoother, more uniform contact surface compared to the laser-machined Mylar gasket that was used in previously published studies. The epoxy layer between the steel spacer and the cover glass adds approximately 40 μm to the height of the chamber. Average chamber height was 138.4 μm +/- 12.1 μm, and the interior volume was approximately 15.65 μL. The average hole diameter was 740.1 +/- 125.1 μm.

### Cell culture and plating for the REEC/4T1 system

Murine mammary triple negative breast cancer 4T1 cells were cultured in complete media (DMEM media, 25 mM glucose, 10% fetal bovine serum (FBS), penicillin, and streptomycin; Quality Biologicals). GFP-expressing 4T1 cells, 4T1-Fluc-Neo/eGFP-Puro cells (GFP-4T1, Imanis Life Sciences), were grown in complete media and selected for using G418 and puromycin. Prior to plating, laser-machined Mylar rims were epoxied on to a 12 well-plate. The Mylar rims ensure that the Mylar clamps of each REEC are able to firmly hold the REEC against the bottom of the well. 4T1 cells were then plated (200,000 cells/mL, 1 mL/well) and incubated at 37°C. Once the 4T1 cells reached 70-80% confluency (^~^24 hours), the media was refreshed and REECs were placed with sterilized forceps and pressed firmly to the bottom of the well. Media in the upper chamber was refreshed every 3-4 days with the least possible disturbance to the lower chamber possible.

### Mathematical model

The REEC/4T1 system was simulated as an annulus containing a uniform density of consumers and used a diffusion-consumption model^20^ to characterize the time-evolution and steady state behavior of oxygen and glucose. For a radially symmetric system, the concentration (C) of a molecular species is described by

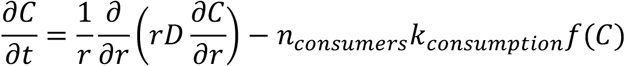

Where *n* = concentration of consumers, *k* = the per-cell consumption rate, and *f*(*C*) is a function of concentration and can be written as

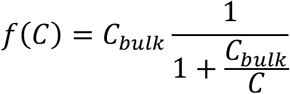

At steady state, a characteristic distance can be defined as

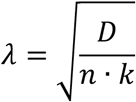

*k*, can be calculated from the maximum per cell molecular consumption rate, *A_max_*, and the concentration of the molecule of interest, *C_bulk_*

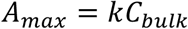

MATLAB’s partial differential equation toolbox was used to model the evolution of oxygen and glucose gradients within the REEC/4T1 system at 37 °C. The system was defined as an annulus with an inner radius, r_1_, of 350 μm and an outer radius, r_2_, of 6,000 μm, corresponding to the radius of the REEC opening and the inner radius of the REEC, respectively. The inner radius was modeled as a source of the diffusing molecules with a constant concentration, C_bulk_.

Thus, the boundary conditions were

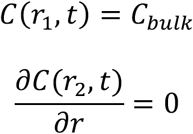

The initial condition corresponded to the moment the chamber was placed in the well:

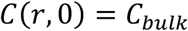

Oxygen diffusion, glucose diffusion, and glucose consumption rate values for 4T1 cells were based on the literature^23–27^. Oxygen consumption rate (OCR) was measured directly via a Seahorse XF96 metabolic analyzer (see below).

The oxygen concentration used in the dynamic model at *r* < 350 μm is 171 μM, not 178 μM which is the value at the interface of the media and the atmosphere. It is somewhat less because the cells immediately below the opening in the REEC consume the oxygen diffusing through the opening in the coverglass. To account for this, we modeled the oxygen concentration in a column of media 250 μm tall (100 μm high REEC and a 150 μm thick coverglass) above a monolayer of 4T1 cells using a model based on Fick’s law^25^. At the top of the column, the media above the REEC coverglass is assumed to be fully oxygenated (178 μM). Using a 4T1 density was 200,000 cells/cm^2^ and the Amax for the 4T1s, the O_2_ concentration at the cell layer was calculated to be 171 μM.

For each molecular species, the simulation was run for an equivalent of 48 hours for each combination of maximum cellular molecular consumption rate, A_max_, and cell density, *n*. Convergence to steady state was defined as a change in RMS difference of less than 0.1 % between successive profiles.

### Oxygen consumption rate measurements

4T1 cells’ OCR was measured using the XF96 Seahorse Metabolic Analyzer (Agilent Technologies, California). 4T1s were plated (1×10^5^ cells) in each well (200 μL) of a Seahorse microplate. The plates were then incubated at 37°C for 2 hours to allow time for the 4T1 cells to adhere. Mitochondrial stress tests were performed per manufacturer’s instructions. The OCR was measured as cells were treated sequentially with oligomycin (inhibitor of complex V in the electron transport chain (ETC)), trifluoromethoxy carbonylcyanide phenylhydrazone (FCCP, Sigma-Aldrich, a depolarizer of the mitochondrial membrane potential), and rotenone and antimycin-A (inhibitors of complex I and III in the ETC, respectively). Basal respiration, ATP-linked respiration, and spare capacity were calculated using the Seahorse software.

### 4T1 mouse mammary tumor model

The NCI-Frederick Animal Facility, accredited by the Association for Accreditation of Laboratory Animal Care International, follows the Public Health Service Policy for the Care and Use of Laboratory Animals. Animal care was provided in accordance with the procedures outlined in the Guide for Care and Use of Laboratory Animals. Protocols for in vivo studies were approved by the Frederick Animal Care and Use Committee (ACUC). Female BALB/c mice obtained from the Frederick Cancer Research and Development Center Animal Production Area were housed five per cage. Eight to ten-week-old female BALB/c mice were subcutaneously injected with 2 × 10^5^ 4T1 cells. The allograpft tumor volume was measured by Vernier caliper and calculated as volume (mm^3^) = (width^2^ × length)/2. When the tumors reached 2000 mm^3^, typically 30 days post injection, the mice were euthanized, tumors were collected for analysis. Tumors were flash frozen in liquid nitrogen and the tissues were cut into 10 μm thick sections by the Pathology/Histotechnology Laboratory at NCI – Frederick.

### Fixation

4T1 cells cultured in 12-well plates were fixed in 4% v/v paraformaldehyde for 15 minutes. Samples were rinsed three times in PBS and then blocked and permeabilized in blocking buffer (3% BSA w/v, 0.3% Triton-X100 in 1X DPBS) for 1 hour.

Fresh frozen sections of 4T1 tumors were fixed in 4% v/v paraformaldehyde for 30 minutes. Samples were rinsed three times in PBS and then blocked and permeabilized in blocking buffer for 1.5 hours.

### Immunofluorescence Staining

After being fixed, blocked, and permeabilized, cultured 4T1 cells were stained with antibodies diluted in blocking buffer. Incubation times, temperatures, dilutions, and secondaries (if necessary) were used as described in Table 2. For overnight incubations, the samples were kept in a humidified chamber. Cells were then washed three times with 1X PBS and stained with DAPI (300nM; ThermoFisher Scientific) for 15 minutes in 1X PBS. Cells were rinsed an additional three times with 1X PBS prior to storage or imaging.

**Table 1.**

| Parameter | Value | Reference |
| --- | --- | --- |
| D ($\mu\text{m}^2/\text{s}$) | | |
| O<sub>2</sub> | 3,370 | <sup>21</sup> |
| Glucose | 616 | <sup>22</sup> |
| A<sub>max</sub><br>(molecules/cell/s) |  |  |
| O<sub>2</sub> | 2.98x10<sup>7</sup> | OCR measured with Seahorse XF mito-stress test with 4T1 cells. |
| Glucose | 6.47x10<sup>7</sup> | <sup>23</sup> |
| C<sub>bulk</sub> |  |  |
| O<sub>2</sub> | 178 $\mu\text{M}$,<br>171 $\mu\text{M}$ | <sup>24, 25</sup> |
| Glucose | 25 mM | DMEM media specification (Quality Biologicals) |

**Table 2:**

| Antibody & fluorophore | Clone | Vendor – Catalog number | Host | Dilution | Time | Temp. |
| --- | --- | --- | --- | --- | --- | --- |
| E-cadherin, direct conjugate (AlexaFluor-488) | 24E10 | Cell Signaling Technologies – 3199S | Rabbit | 1:100 | O/N | 4°C |
| Vimentin, direct conjugate (AlexaFluor-555) | D21H3 | Cell Signaling Technologies – 9855S | Rabbit | 1:50 | O/N | 4°C |
| DAPI | NA | ThermoFisher | NA | 1:1000 | 15 min | RT |
| Nos2, direct conjugate (AlexaFluor-568) | EPR16635 | AbCam – 209595 | Rabbit | 1:100 | 1 h or O/N | RT or 4°C |
| Cox2, direct conjugate (AlexaFluor-488) | D5H5 | Cell Signaling Technologies – 13596S | Rabbit | 1:100 | 1 h or O/N | RT or 4°C |
| HIF1α, direct conjugate (AlexaFluor-647) | EPR16897 | AbCam – 208420 | Rabbit | 1:100 | O/N | 4°C |
| ATP5A1 (Mitochondrial ATPase, Primary) | NA | AbClonal – A5884 | Rabbit | 1:100 | O/N | 4°C |
| (Anti-rabbit CF-640R secondary for ATPase) | NA | Biotium – 20178 | Donkey | 1:100 | O/N | 4°C |
| α-Smooth Muscle Actin, direct conjugate (eFluor-488), eBioscience | 1A4 | ThermoFisher - 41-9760-82 | Mouse | 1:400 | O/N | 4°C |

After being fixed, blocked, and permeabilized, fresh frozen sections of 4T1 tumors were stained overnight at 4°C with antibodies diluted in blocking buffer, as described in Table 2. Samples were rinsed three times with PBS, stained with DAPI (300nM) for 30 minutes, rinsed again, and sealed for imaging on the Nikon Eclipse Ti widefield fluorescence microscope.

### Multiplexed Immunofluorescence

After imaging with appropriate filter sets for the directly conjugated antibodies, cell and tissue samples were quenched^26, 27^ using a solution comprised of 1 part hydrogen peroxide, 1 part sodium bicarbonate (pH 10) and 3 parts water for 15 minutes at room temperature. Samples were then washed three times with 1X PBS and imaged to ensure quenching. Samples were re-blocked with blocking buffer as described above and re-stained with a new set of directly conjugated antibodies. The quenching cycle can be repeated for at least four rounds of staining without noted damage. DAPI does not quench and can be used to register images for processing. Imaging was performed on the Nikon Eclipse Ti widefield fluorescence microscope.

### Stimulation of ANA-1 macrophages

Murine ANA-1 macrophage cell line was established by infection of normal bone marrow from C57BL/6 mice with J2 recombinant virus^28, 29^. They were cultured in complete media. ANA-1s were stimulated via treatment with IFNγ (100 U/mL) and LPS (20 ng/mL) for 18 to 24 hours. The media was then removed and replaced with fresh stimulation media and subsequent experiments were performed immediately.

### ANA-1 injection into 4T1 REECs

In order to allow time for disks to form, REECs were placed on a 70-80% confluent monolayer of 4T1-GFP-luc cells, using the method described above, 3 days prior to ANA-1 injection. ANA-1 cells were stimulated with IFNγ and LPS, as described above, 1 day prior to ANA-1 injection. On the day of ANA-1 injection, ANA-1 cells were incubated in serum-free media with CellTracker Red CMTPX Dye (5 μM, ThermoFisher) for at 37 °C for 30 minutes. The ANA-1 cells were then spun down and diluted in complete media (+/- IFNγ and LPS) to ^~^10,000,000 cells/mL. Immediately prior to injection, the media in the upper chamber of the REECs was refreshed with complete media (+/- IFNγ and LPS). The concentrated ANA-1 solution (10 μL, containing ^~^100,000 ANA-1 cells) was injected through the opening, directly into the lower chamber of the REEC. For controls, complete media (10 μL, +/- IFNγ and LPS) containing no ANA-1 cells was injected instead. Images were taken every 2 hours for 48 hours on the Nikon Eclipse Ti widefield fluorescence microscope with a 4X dry objective and using a live cell heated stage.

### Extracellular O_2_ concentration quantification in REECs via O_2_ sensor foils

To quantify the spatial variation of dissolved [O_2_] in the media across the REEC we used O_2_ sensor foils (PreSens Precision Sensing GmbH, Germany) -- 100 μm thick hydrogels impregnated with particles with [O_2_]-dependent luminescence. The ratio of the red to blue luminescence correlates with dissolved O_2_ concentration. The O_2_ sensor foils were glued to the inside top surface of the REEC (made with 200 μm thick stainless-steel washers to maintain a 100 μm chamber height above cells) and a hole was drilled through both the glass and the foil. The response of the sensor foils to [O_2_] was calibrated prior to placement on cells using glucose oxidase/catalase solutions of known [O_2_] using the Nikon microscope with the 4X objective lens, using the TRITC and DAPI emission filters in rapid succession and exciting the foil with a 395 nm LED lamp. The [O_2_] in the glucose oxidase/catalase solution was measured using a Piccolo oxygen sensor system (PyroScience GmbH, Germany). To measure the [O_2_] dynamics across the REEC with 4T1 cells in phenol red-free media (1 mL), images of the sensor foil were collected every 15 minutes using the 4X objective lens. A flat-field correction was applied to the images, and the red/blue intensity ratio as a function of distance from the center of the hole was converted via the calibration curve to dissolved [O_2_] values (Supplemental Figure 3).

### Extracellular hypoxia gradient dynamics in REECs via Image-iT Green Hypoxia Reagent

To measure the development of gradients of intracellular hypoxia of live cells in the REECs, cells were incubated with Image-iT Green Hypoxia Reagent (IGHR) (10 μM; Invitrogen) at 37 °C for 30 minutes prior to chamber placement. After chamber placement, 4T1 cells were imaged using standard FITC excitation and emission filters at 4X or 20X magnification on a Nikon Eclipse Ti widefield fluorescence microscope. For controls, IGHR treated 4T1 cells were placed in incubators set to 0.1%, 1%, or 5% O_2_ or in a standard incubator (^~^20%) for 2 hours. Cells were immediately imaged (Supplemental Figure 3).

### Intracellular Hypoxia quantification in REECs

To measure levels of intracellular hypoxia in cell disks, media in the upper chamber was replaced with complete media supplemented with pimonidazole (200 μM; Hypoxyprobe, Inc., Massachusetts) after disk formation. In hypoxic cells, pimonidazole is reduced and forms adducts with thiol groups. After sufficient time for the pimonidazole to diffuse through the chamber and be taken up by cells (^~^6 hours), chambers and media were removed, and the disks were immediately fixed, blocked, and permeabilized as described above. A monoclonal antibody specific to pimonidazole adducts and conjugated with a fluorescein probe (Hypoxyprobe-Green Kit (FITC-Mab); Hypoxyprobe, Inc.) was applied for 1 hour at room temperature, or overnight at 4 °C. Samples were rinsed three times with 1X PBS and imaged using standard FITC excitation and emission filters at 20X magnification on the Nikon Eclipse Ti widefield fluorescence microscope. For controls, cultured 4T1 cells were treated with complete media which had been supplemented with pimonidazole and deoxygenated (<1% O_2_) with glucose oxidase and catalase. HIF1α expression was determined by immunofluorescence and correlated with pimonidazole adduct formation (Figure 1h).

### Glucose Gradient dynamics in REECs

2 hours after chamber placement, (2-(N-(7-Nitrobenz-2-oxa-1,3-diazol-4-yl)Amino)-2-Deoxyglucose (2-NBDG) (100 μM; ThermoFisher Scientific) with Hoechst (1 μg/mL) in low-glucose media (5 mM glucose, 10% FBS) was diffused into the chamber for 30 minutes at 37 °C. This dye was often combined with tetramethylrhodamine, ethyl ester (TMRE) (1 nM; ThermoFisher Scientific) in order to obtain simultaneous metabolic measurements. Chambers were removed, rinsed three times in 1X PBS, and imaged using standard excitation and emission filters at 20X on the Nikon Eclipse Ti widefield fluorescence microscope. Samples could be fixed. For measurements in the disk, similar methods were used 7 days after chamber placement on a Zeiss 710 Laser Scanning Confocal Microscope with a 10X dry objective.

### Live/Dead Staining

4-6 days after chamber placement, media was replaced with media containing the live/dead cell stains ethidium homodimer (2 μM), calcein AM (3 μM), and Hoechst (1 μg/mL; ThermoFisher) for 30 minutes. Cells were then imaged on the Nikon Eclipse Ti widefield fluorescence microscope using a live cell heated stage.

### Celigo Image Acquisition and Analysis

Cells were cultured in 12-well plates with chambers for up to 21 days with weekly media replacement. The plate was scanned using the Cell Counting feature in the Celigo Imaging Cytometer (Nexelom Biosciences) with the brightfield algorithm to detect cells. Cells were segmented using the built-in software.

### Nos2 and Cox Inhibitor treatments

Immediately prior to chamber placement, the media in each well was replaced with treatment media: complete media supplemented with either the Cox inhibitor indomethacin (100 μM) or the Nos2 inhibitor aminoguanidine (1 mM). Cells were maintained in the REECs for 3-7 days using treatment media before fixation and immunofluorescence staining. For controls, standard complete media was used.

### Nitric Oxide Production in REECs

4-Amino-5-Methylamino-2’,7’-Difluorofluorescein Diacetate (DAF) (ThermoFisher) was used to measure and spatially resolve nitric oxide (NO) production in disks. 7 days after chamber placement, wells were washed three times in 1X PBS to remove all phenol red and BSA, which interfere with DAF fluorescence. To ensure that the lower compartment of the chamber was adequately washed, the 1X PBS was gently pipetted up and down directly over the opening. The 4T1 cells were then incubated in phenol-red free, serum free media with DAF (10 μM) and Hoechst (μg/mL) at 37 °C for 45 minutes. This dye was often combined with TMRE in order to obtain simultaneous measurements of mitochondrial membrane polarization state. Samples were then immediately imaged on a Zeiss 710 Laser Scanning Confocal Microscope with a 10X dry objective using standard FITC emission and excitation filters.

### Metabolic Gradient Quantification in REECs

Cells were incubated in phenol red free media with TMRE at 37°C for 20 minutes for cells in standard culture, or 45 minutes for cells in a REEC. TMRE was used simultaneously with 2-NBDG or DAF Diacetate. Cells were imaged with or without chamber removal, depending on the experiment. For controls, the electron transport chain was inhibited with FCCP or antimycin A and rotenone. These treatments caused the TMRE signal to decrease significantly within 15 minutes. TMRE was imaged on a Zeiss 710 Laser Scanning Confocal Microscope with a 10X dry objective using Texas Red excitation and emission filters.

To confirm that mitochondrial mass was consistent across the disk, disks were then fixed, blocked, and permeabilized, as described above. The disks were stained with an ATP-synthase antibody (Abclonal, A5884, Rabbit) diluted 1:100, and incubated at 4°C overnight. Cells were rinsed and incubated with an anti-rabbit secondary at room temperature for 1 hour and then imaged on a Zeiss 710 Laser Scanning Confocal Microscope with a 10X dry objective.

### Spheroid Growth, Clearing, and Imaging

A spheroid formation assay was performed in ultra-low attachment round bottom 96 well plates (Nexcelom, Lawrence, MA, USA). 4T1 cells or 4T1-GFP-luc cells (for co-culture experiments) were plated in each well (200 μL, 6 x 10^3^/mL) with serum-free DMEM supplemented with basic Fibroblast Growth Factor (20 ng/mL) and B-27 supplement (1:50; Thermo Fisher Scientific). Media was supplemented on Day 4. Monoculture spheroids were fixed, cleared, immunolabeled with antibodies and imaged on Day 7 (see below). For coculture experiments, ANA-1 macrophages were stimulated for 18 hours followed by treatment with CellTracker Red CMTPX dye (10μM) at 37°C for 45 minutes and were then added to the spheroids (2 x 10^5^ cells/well). The ANA-1-spheroid cocultures were fixed, cleared, immunocytochemically stained with antibodies and imaged on Day 7 (see below).

Spheroids were cleared for imaging using the Ce3D method^30^. Briefly, spheroids were fixed in 4% v/v paraformaldehyde containing 0.5% Triton-X100. Spheroids were blocked at 37°C for 36 hours in a humidified environment. Spheroids were then stained with directly conjugated antibodies for proteins of interest (HIF1α, Nos2, Cox2) at 4 °C for 36 hours. Spheroids were stained with DAPI (300 nM) for 30 minutes and then rinsed three times with 1X PBS. The spheroids were then embedded in 1.5% low-melt agarose. Samples were placed in clearing solution (0.1% v/v Triton-X100, 13% N-methylacetamide, 66% w/v Nycodenz AG) at room temperature for 4 hours at a 1:4 agarose:clearing solution ratio. After 4 hours, the clearing solution was replaced with the equivalent volume of fresh clearing solution and left overnight. The cleared spheroids were imaged on a Leica TCS SP8 Laser Scanning Confocal microscope at 20X with oil to match the index of refraction of the clearing solution (slightly higher than 1.5). Z-stacks were taken through the center of the spheroids.

### Image Processing and Data Analysis

Images taken on the Nikon Eclipse Ti widefield fluorescence microscope were typically stitched and background corrected using a rolling ball technique in Nikon Elements software, then processed in Imaris (Bitplane). Image brightness and contrast were adjusted to optimize the visual dynamic range for display. Imaris was used for cell segmentation and to extract position, fluorescence intensity, and geometrical statistics on each cell. Custom R scripts were used to process the output of Imaris statistics, including average cell intensity for each channel and position. For cell disks, cells were binned into annuli every 50 μm from the center, and the mean fluorescence intensity (MFI) per cell in that bin was calculated. Disks that could not be segmented were analyzed in FIJI using the Radial Profile plug-in, which averages the intensity value of all pixels at each radius from a fixed point. For the spheroid and tissue images, custom MATLAB functions were used to calculate fluorescence intensity and depth (spheroid) or position (tissue) for each pixel and averaged as above. For disks and spheroids intensity profiles, R scripts were used to average those radial mean values within 50 μm wide annuli. For cell disks, radial values were normalized to disk radius, and MFI were normalized to the value at r = 0 for that disk or the control group disk. For the NBDG/TMRE data, MFI were normalized to the maximum MFI value for each profile, and the normalized curves were averaged. For spheroids, depth values were normalized to the maximum depth of each spheroid (such that 0 represents the surface and 1 is the maximum depth), and MFI were normalized to the value at the spheroid surface. For tissue, MFI was normalized to the first position in the profile. All data are presented as mean +/- SEM unless otherwise noted. Statistical significance, determined using Welch’s two-tailed t-test, and Pearson correlation coefficients were calculated in GraphPad Prism and Microsoft Excel.

## Supporting information

Supplemental Figures and Protocol

## Data Availability

The data that support the findings of this study are available from the corresponding author upon request.

## Code Availability

The code supporting the plots and other findings in the manuscript are available from the corresponding author upon request.

## Acknowledgments

This project has been funded in whole or in part with Federal funds from the National Cancer Institute, National Institutes of Health, under Contract No. 75N91019D00024 and by the Intramural Program of the NIH, NCI, Center for Cancer Research. The content of this publication does not necessarily reflect the views or policies of the Department of Health and Human Services, nor does mention of trade names, commercial products, or organizations imply endorsement by the U.S. Government. NCI-Frederick is accredited by AAALAC International and follows the Public Health Service Policy for the Care and Use of Laboratory Animals. Animal care was provided in accordance with the procedures outlined in the “Guide for Care and Use of Laboratory Animals” (National Research Council; 2011; National Academy Press; Washington, D.C.). In addition, we would like to thank Dan McVicar for the generous use of his Seahorse XF Analyzer.

## Author contributions

WFH, SJL, and DAW conceived the project, and ACG, SJF, VS, and WFH designed the experiments. ACG and SJF constructed the REECs, performed all experiments with the chambers, wrote analysis software, and processed and analyzed the images. VS grew the spheroids and performed the mouse experiments and assisted ACG with the Seahorse measurements. ACG imaged the spheroids and tissue samples. DAS assisted with multiplexed immunofluorescence imaging and image analysis. WFH implemented the mathematical model. ACG, SJF, and WFH wrote the manuscript, and all authors critically reviewed the manuscript.

## References

1. Petrova, V., Annicchiarico-Petruzzelli, M., Melino, G. & Amelio, I. The hypoxic tumour microenvironment. Oncogenesis 7 (2018).

2. Byrne, M.B., Leslie, M.T., Gaskins, H.R. & Kenis, P.J.A. Methods to study the tumor microenvironment under controlled oxygen conditions. Trends in Biotechnology 32, 556–563 (2014).

3. Keenan, T.M. & Folch, A. Biomolecular gradients in cell culture systems. Lab on a Chip 8, 34–57 (2007).

4. Friedrich, J., Seidel, C., Ebner, R. & Kunz-Schughart, L.A. Spheroid-based drug screen: Considerations and practical approach. Nature Protocols (2009).

5. Nath, S. & Devi, G.R. Three-dimensional culture systems in cancer research: Focus on tumor spheroid model. Pharmacology and Therapeutics 163, 94–108 (2016).

6. Hoh, J.H., Werbin, J.L. & Heinz, W.F. Restricted exchange microenvironments for cell culture. BioTechniques 64, 1–6 (2018).

7. Pulaski, B.A. & Ostrand-Rosenberg, S. Mouse 4T1 Breast Tumor Model. Current Protocols in Immunology 39, 20.22.21–20.22.16 (2000).

8. Noy, R. & Pollard, Jeffrey W. Tumor-Associated Macrophages: From Mechanisms to Therapy. Immunity 41, 49–61 (2014).

9. Lehmann, S. et al. Hypoxia Induces a HIF-1-Dependent Transition from Collective-to-Amoeboid Dissemination in Epithelial Cancer Cells. Current Biology (2017).

10. Nieto, M.A., Huang, R.Y.-j., Jackson, R.A. & Thiery, J.P. Review EMT : 2016. Cell 166, 21–45 (2016).

11. Basudhar, D. et al. Coexpression of NOS2 and COX2 accelerates tumor growth and reduces survival in estrogen receptor-negative breast cancer. Proceedings of the National Academy of Sciences 114, 13030–13035 (2017).

12. Somasundaram, V. et al. Molecular Mechanisms of Nitric Oxide in Cancer Progression, Signal Transduction, and Metabolism. Antioxidants & Redox Signaling 30, 1124–1143 (2018).

13. Heinecke, J.L. et al. Tumor microenvironment-based feed-forward regulation of NOS2 in breast cancer progression. Proc Natl Acad Sci U S A 111, 6323–6328 (2014).

14. Basudhar, D. et al. Understanding the tumour micro-environment communication network from an NOS2/COX2 perspective. Br J Pharmacol 176, 155–176 (2019).

15. Somasundaram, V. et al. Inducible nitric oxide synthase-derived extracellular nitric oxide flux regulates proinflammatory responses at the single cell level. Redox Biology 28, 101354 (2020).

16. Olson, N. & Van Der Vliet, A. Interactions between nitric oxide and hypoxia-inducible factor signaling pathways in inflammatory disease. Nitric Oxide - Biology and Chemistry 25, 125–137 (2011).

17. Cyr, A. et al. Endotoxin Engages Mitochondrial Quality Control via an iNOS-Reactive Oxygen Species Signaling Pathway in Hepatocytes. Oxid Med Cell Longev 2019, 4745067 (2019).

18. Vitale, I., Manic, G., Coussens, L.M., Kroemer, G. & Galluzzi, L. Macrophages and Metabolism in the Tumor Microenvironment. Cell Metab 30, 36–50 (2019).

19. Jeong, S.K. et al. Tumor associated macrophages provide the survival resistance of tumor cells to hypoxic microenvironmental condition through IL-6 receptor-mediated signals. Immunobiology 222, 55–65 (2017).

20. Oyler-Yaniv, A. et al. A Tunable Diffusion-Consumption Mechanism of Cytokine Propagation Enables Plasticity in Cell-to-Cell Communication in the Immune System. Immunity 46, 609–620 (2017).

21. Pettersen, E.O., Larsen, L.H., Ramsing, N.B. & Ebbesen, P. Pericellular oxygen depletion during ordinary tissue culturing measured with oxygen microsensors. Cell Proliferation 38, 257–267 (2005).

22. Suhaimi, H., Wang, S. & Das, D.B. Glucose diffusivity in cell culture medium. Chemical Engineering Journal 269, 323–327 (2015).

23. Simões, R.V. et al. Metabolic Plasticity of Metastatic Breast Cancer Cells: Adaptation to Changes in the Microenvironment. Neoplasia 17, 671–684 (2015).

24. Wenger, R., Kurtcuoglu, V., Scholz, C., Marti, H. & Hoogewijs, D. Frequently asked questions in hypoxia research. Hypoxia 3, 35 (2015).

25. Al-Ani, A. et al. Oxygenation in cell culture: Critical parameters for reproducibility are routinely not reported. PLoS One 13, e0204269 (2018).

26. Gerdes, M.J. et al. Highly multiplexed single-cell analysis of formalin-fixed, paraffin-embedded cancer tissue. Proceedings of the National Academy of Sciences 110, 11982–11987 (2013).

27. Lin, J.R., Fallahi-Sichani, M., Chen, J.Y. & Sorger, P.K. Cyclic Immunofluorescence (CycIF), A Highly Multiplexed Method for Single-cell Imaging. Current protocols in chemical biology 8, 251–264 (2016).

28. Blasi, E., Radzioch, D., Durum, S.K. & Varesio, L. A murine macrophage cell line, immortalized by v-raf and v-myc oncogenes, exhibits normal macrophage functions. Eur J Immunol 17, 1491–1498 (1987).

29. Blasi, E. et al. Selective immortalization of murine macrophages from fresh bone marrow by a raf/myc recombinant murine retrovirus. Nature 318, 667–670 (1985).

30. Li, W., Germain, R.N. & Gerner, M.Y. Multiplex, quantitative cellular analysis in large tissue volumes with clearing-enhanced 3D microscopy (C e 3D). Proceedings of the National Academy of Sciences (2017).

