## Supplemental Figures and Protocol for "An *in vitro* tumorigenesis model based on live cell-generated oxygen and nutrient gradients"

### Supplemental Material for:
### An *in vitro* tumorigenesis model based on live cell-generated oxygen and nutrient gradients

Anne C. Gilmore, Sarah J. Flaherty, Veena Somasundaram, David A. Scheiblin[3], Stephen J. Lockett, David A. Wink, William F. Heinz*

**This PDF file includes:**

Supplemental Figures
Supplemental Protocol

**Other Supplementary Materials for this manuscript include:**

Supplemental Video 1

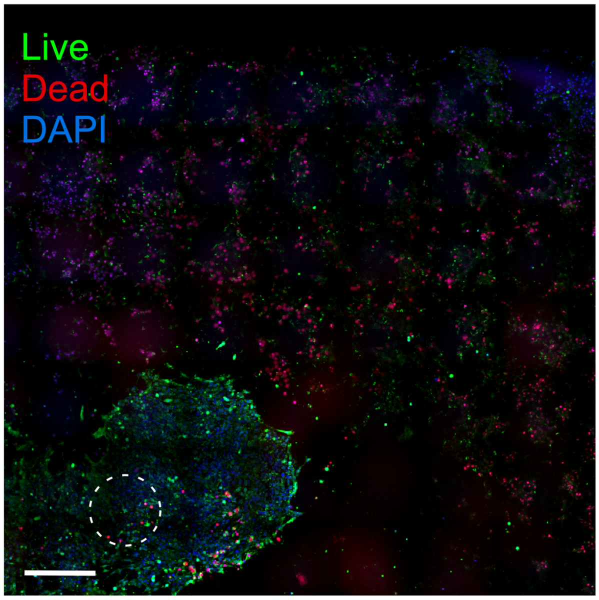


**Supplemental Figure 1. Cell viability across the REEC.** Live/dead staining shows necrotic zone beyond the cell disk live/dead staining in 4-day old 4T1 disks. Live cells are stained by calcein AM (green). Dead cells are stained by ethidium homodimer (red) (scale bar = 500 µm).

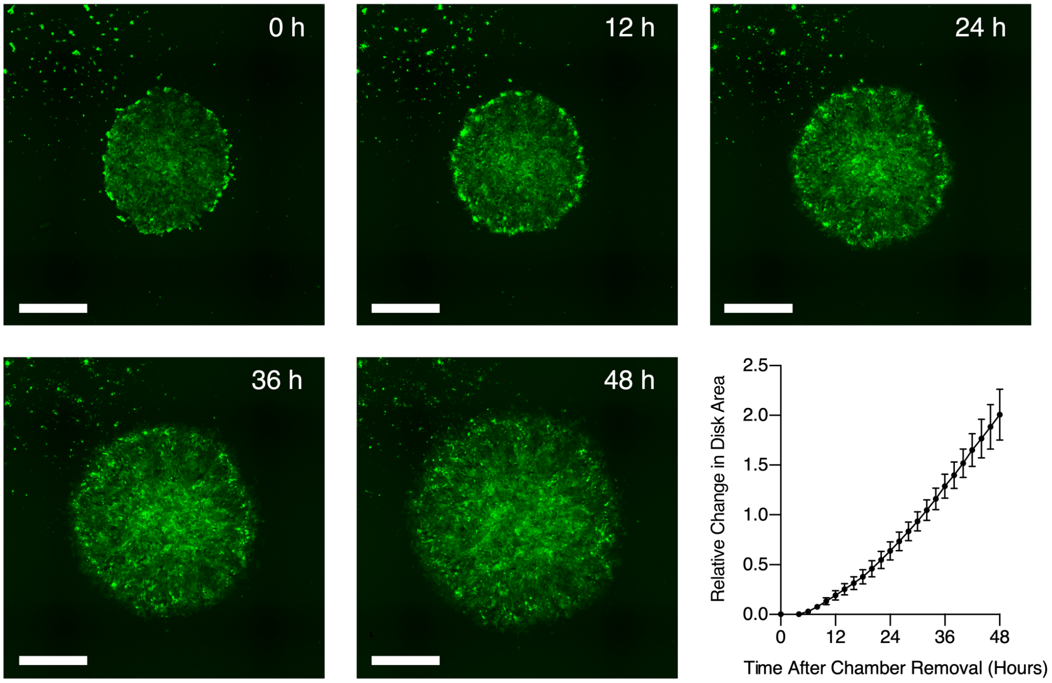


**Supplemental Figure 2. 4T1 proliferation is restored after the REEC is removed and normoxia is established across the disk.** After disk formation (92 hours after chamber placement), the REEC was removed and 4T1 growth was monitored over 48 hours (scale bar = 200 µm).

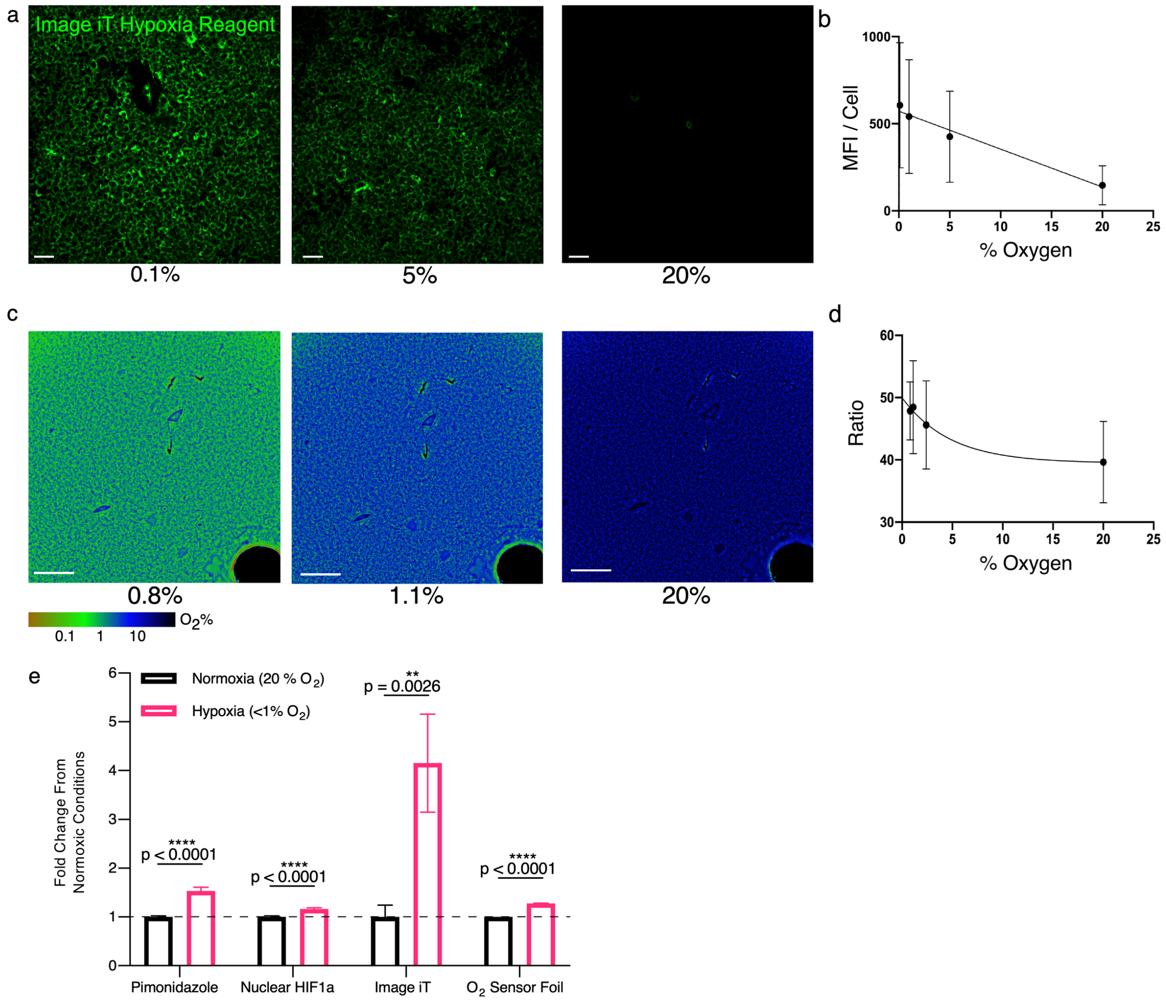


**Supplemental Figure 3. Oxygen sensing techniques.** Multiple complementary techniques were used to measure $[O_2]$ and cellular hypoxia in the REECs including a direct measure of dissolved $[O_2]$ and live-cell and end-point probes of intracellular hypoxia. All methods respond to $[O_2] < 1\%$ in the REEC system.

a. Widefield images of Image IT Green Hypoxia Reagent (IGHR) response to hypoxia in controlled, incubator settings (scale bar = 50 µm).
b. Linear calibration curve of IGHR mean fluorescence intensity per cell to measured oxygen concentration (0.1%, N = 13,214; 1%, N = 27,628; 5%, N = 8,815, 20%, N = 10. N = cell number). Note that at 20% $O_2$, few cells had measurable fluorescence.
c. Ratiometric images of PreSens $O_2$-sensing foil response to $[O_2]$ levels in glucose oxidase solutions (scale bar = 500 µm).
d. Exponential-decay calibration curve of $O_2$-sensing foil ratiometric response in control glucose-oxidase solutions (N = 4).
e. Quantification of fold change in fluorescent signal compared to normoxic conditions of pimonidazole, nuclear-localized HIF1$\alpha$, IGHR, and PreSens $O_2$-sensing foils $O_2$ in response to low oxygen (<1%) (N > 100 cells or pixels, for $O_2$ sensors).

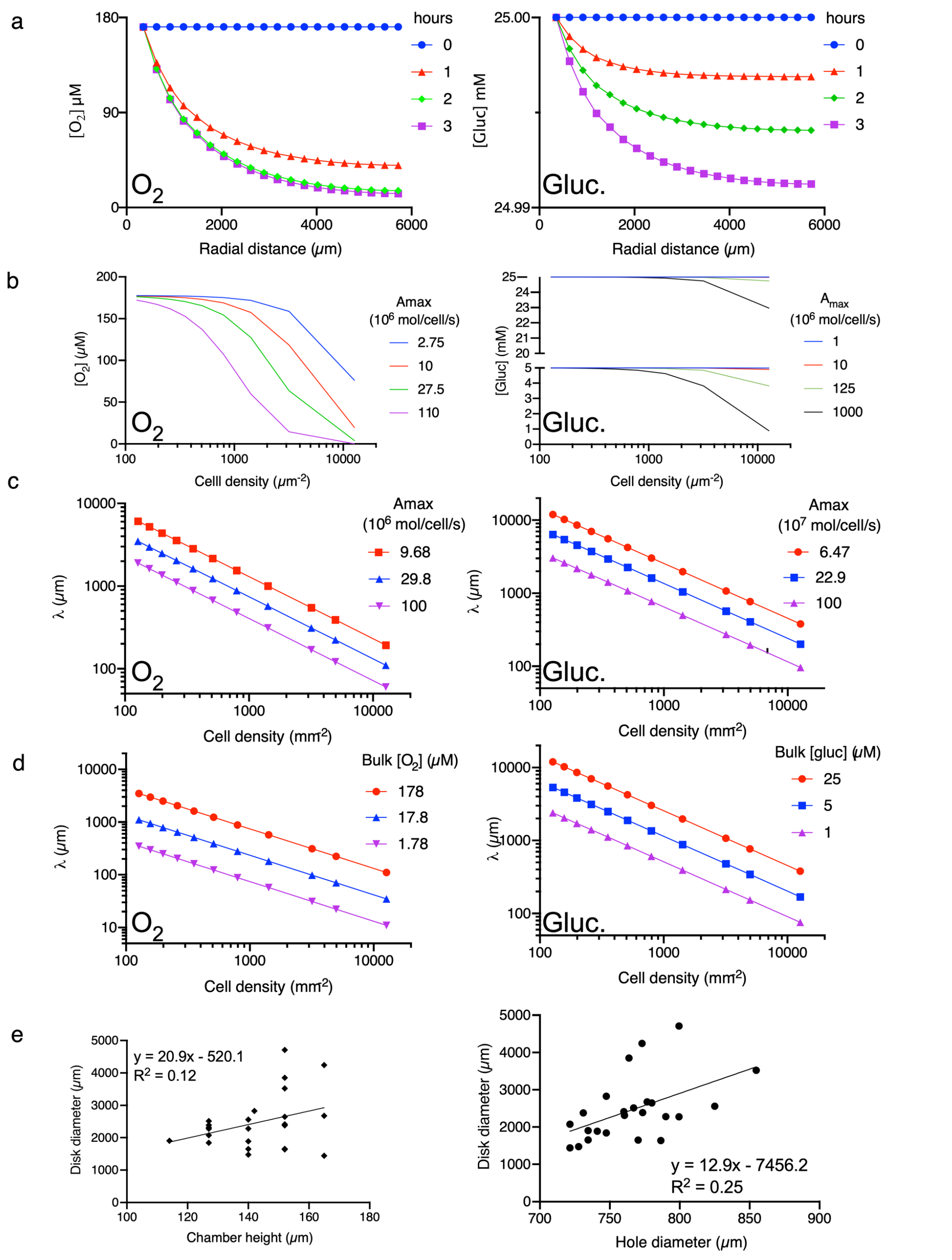


**Supplemental Figure 4. Diffusion-consumption 2D in silico model captures the gradient dynamics and steady state of the 4T1-REEC system.**

a. Time evolution of the radial profiles of the $O_2$ (left) and glucose (right) concentrations (30-minute time steps) calculated with the chamber dimensions (hole diameter = 700 μm), 4T1 consumption rates ($2.98 \times 10^7$ and $6.47 \times 10^7$ molecules/cell/s for $O_2$ and glucose, respectively), cell densities (cell diameter = 13.5 μm), and bulk concentrations used *in vitro* ($[O_2]$ = 171 μM, [Gluc.] = 25 mM). The $[O_2]$ decreases to a stable gradient within 2 h. The glucose concentration changes little because the bulk concentration, 25 mM, is in excess of the cells' metabolic demands.

b. $O_2$ (left) and glucose (right) concentrations as a function of cell density and maximum consumption rate in normoxic nutrient-rich media, $A_{max}$, after 48 h. Glucose concentration curves are shown for bulk concentrations of 25 mM (top) and 5 mM (bottom).
c. Characteristic steady-state distance, $\lambda = \sqrt{DC_{bulk}/nA_{max}}$, as a function of cell density, *n*, and maximum consumption rate, $A_{max}$. *D* is diffusion coefficient. λ is related to the stable cell disk radius. This estimation is independent of REEC geometry. Bulk concentrations ($C_{bulk}$) were 178 µM and 25 mM for $O_2$ (left) and glucose (right), respectively.
d. Characteristic steady-state distance as a function of cell density and bulk concentration. Here, $A_{max}$ values were $2.98x10^7$ and $6.47x10^7$ molecules/cell/s for $O_2$ (left) and glucose (right), respectively.
e. Chamber height and hole diameter dimensions affect final cell disk size.

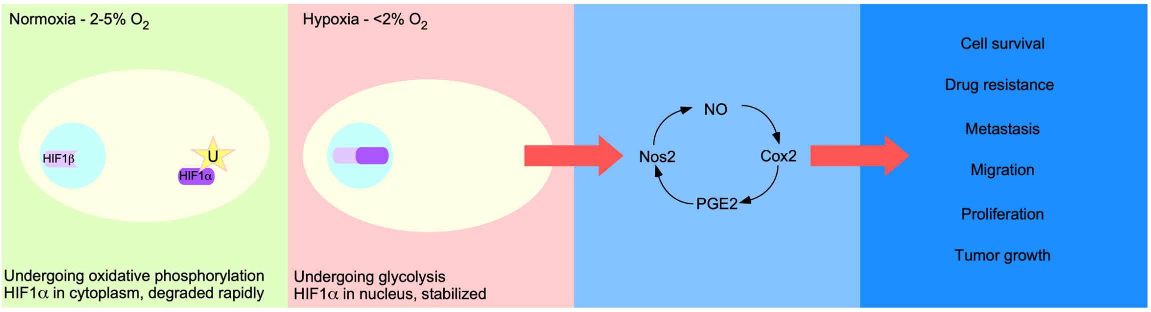


**Supplemental Figure 5. Cellular Response to Hypoxia.** In normoxic conditions HIF1α is rapidly ubiquinated and degraded in the cytoplasm, and HIF1β localizes to the nucleus (left). In hypoxic conditions HIF1α localizes to the nucleus to form a stable dimer with HIF1β (left center). This leads to cell survival and a tumorigenic phenotype, as well as upregulation of the inflammatory protein Nos2 (right center). Nos2 and Cox2 engage in a feedforward loop via nitric oxide (NO) and prostaglandin E2 (PGE2) signaling. Nos2 expression can promote a pro-tumor phenotype (right).

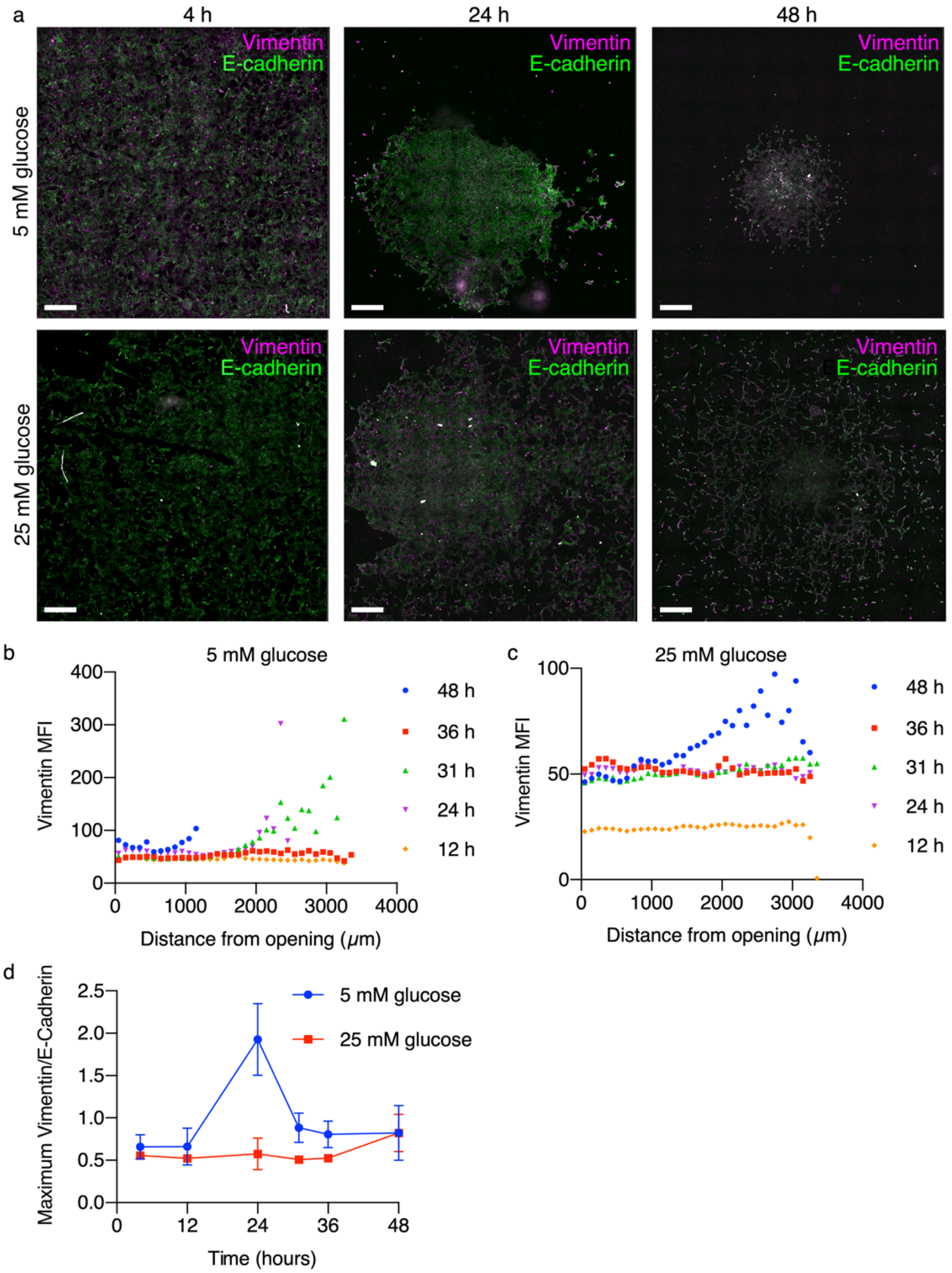


**Supplemental Figure 6. Epithelial mesenchymal transition (EMT) marker expression depends on glucose concentration and time.**

a. Widefield images of vimentin and e-cadherin multiplexed immunofluorescence staining of 4T1 cells cultured in REECs in low (5 mM) or high (25 mM) glucose media at various timepoints after chamber placement (scale bar = 500 µm).
b. Expression of vimentin in low glucose (5 mM) media as a function of distance from opening at different time points (N = 1 per timepoint). Vimentin expression is transiently increased in the hypoxic regions of the disk from 24 to 31 hours during disk formation.

c. Expression of vimentin in high glucose (25mM) media as a function of distance from opening at different time points (N = 1 per timepoint). The 12 h timepoint is low because vimentin is an endpoint EMT marker, and at 12 h EMT has not yet begun. At later times the vimentin expression is elevated. We hypothesize that the levels will return to these low levels once disk formation is complete.
d. Ratio of maximum vimentin to e-cadherin expression in disks for different glucose concentrations over time (N = 3 disks per time point per concentration). The e-cadherin levels did not vary appreciably across the REEC over time. In the 5 mM case, after peaking during disk formation, the ratio returns to the initial value in stable disks.

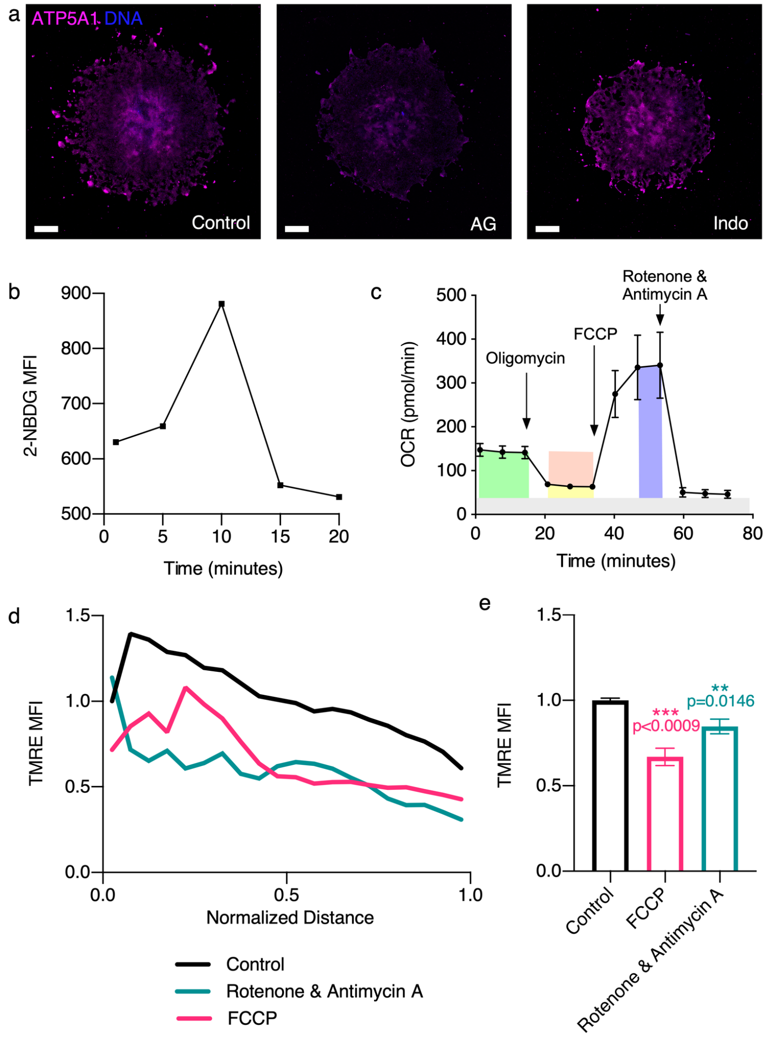


**Supplemental Figure 7. Mitochondrial controls.**

a. Widefield images of ATP5A, a mitochondrial ATPase used here as a measure of mitochondrial mass, immunofluorescence staining in 7-day old 4T1 disks, including a control disk (left), a disk treated with aminoguanidine (middle), and a disk treated with indomethacin (right) (scale bar = 500 µm.)
b. 2-NBDG uptake in 4T1 cells over time in standard culture vessel.
c. 4T1 oxygen consumption rate (OCR) measured by Seahorse Mito Stress test (N = 4 wells). Yellow represents basal respiration, green represents ATP-linked respiration, orange represents proton leak, purple represents maximal respiration, and grey represents non-mitochondrial contributions to OCR, respectively.
d. Quantification of TMRE response (mean fluorescence intensity normalized to first point of the untreated control) across 4T1 disks before and after treatment with rotenone and antimycin-A or FCCP. Images were collected 20 minutes after treatments.
e. Quantification of TMRE response (normalized to the untreated control) before and after treatment with antimycin-A and rotenone or FCCP before and after treatments in standard cell culture conditions.

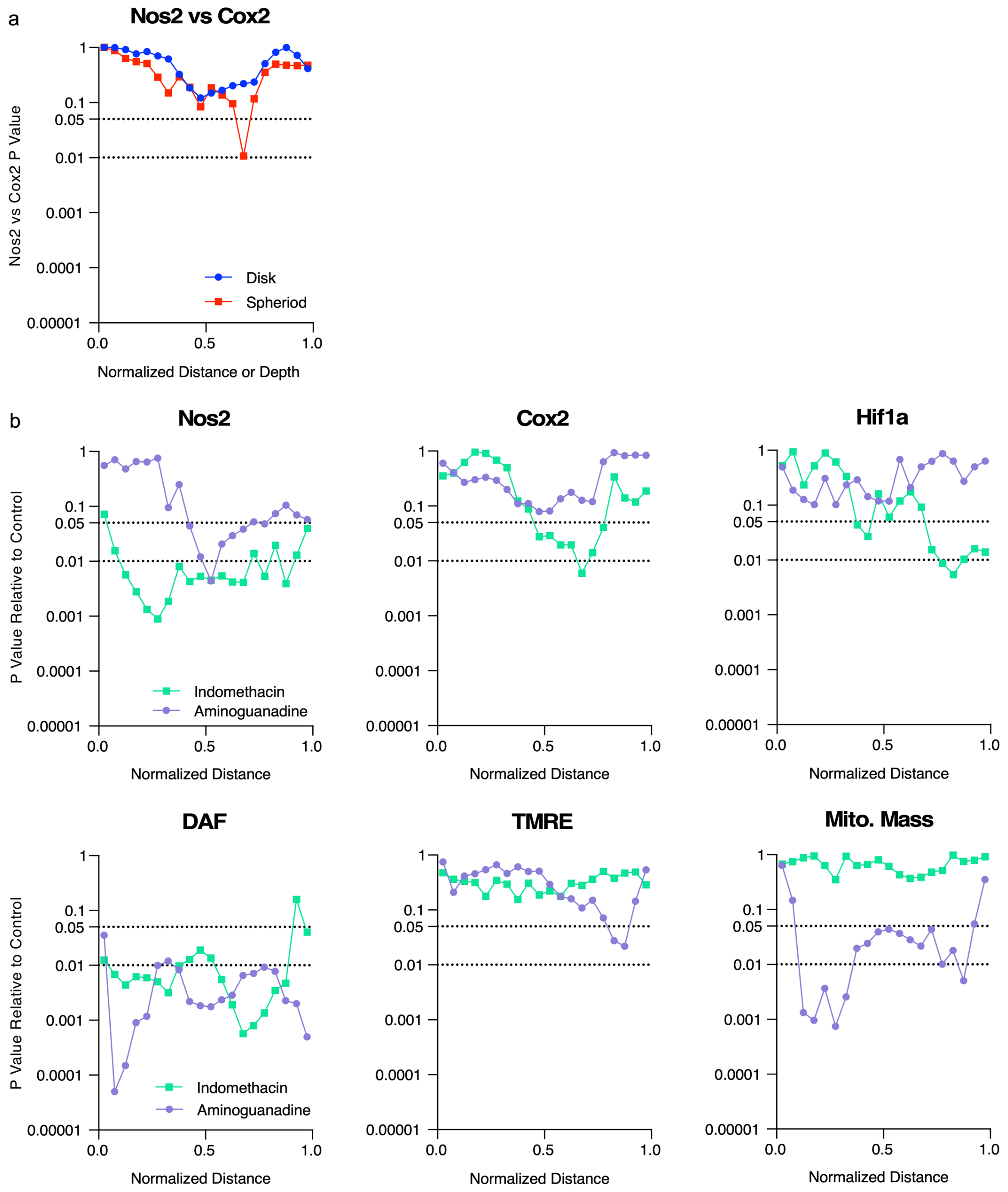


**Supplemental Figure 8. Statistics supplement for Figures 3 and 4.** P-Values were calculated at the normalized distances or depths corresponding to data points in Figures 3 and 4.

a. P-values quantifying the differences between the Nos2 and Cox2 MFI/cell values in the control disks and spheroids that were presented in Fig. 3. Note that the Nos2 and Cox2 signals are most different from one another in the mid-hypoxic region of both disks and spheroids (Normalized distance or depth between 0.5 and 0.75).

b. P-values quantifying the differences between the MFI/cell values observed in the control disks and the MFI/cell values observed in the aminoguanidine- or indomethacin-treated disks that were presented in Fig. 4. The p-values were calculated using the MFI/cell of Nos2, Cox2, HIF1$\alpha$, DAF, TMRE, and mitochondrial mass (measured by ATP5A).

**“**Normalized Distance” refers to the relative distance from the center to the edge of a disk. “Normalized Depth” refers to relative distance from the surface to the center of a spheroid. MFI = mean fluorescence intensity normalized to the first point.

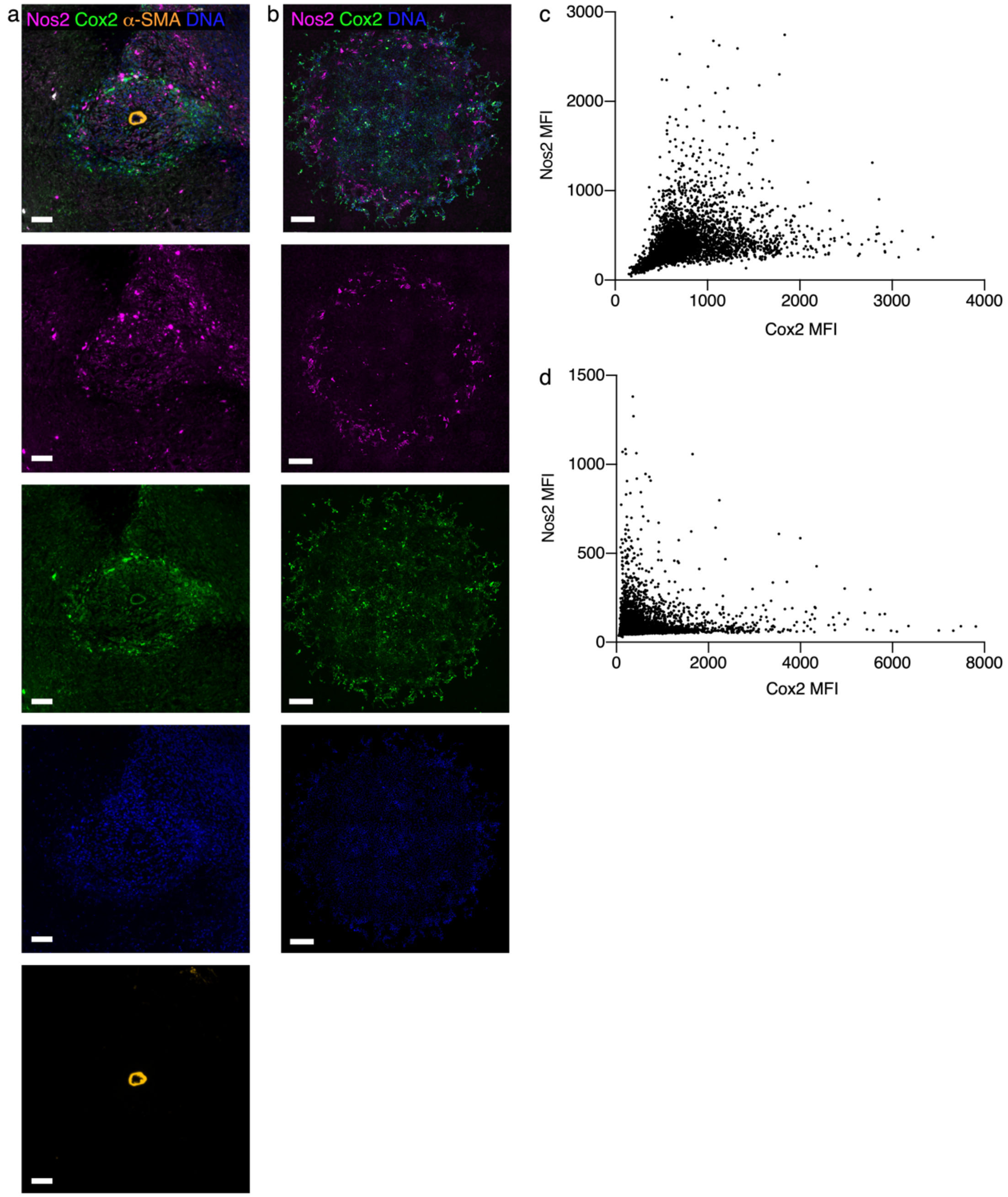


**Supplemental Figure 9. Nos2 and Cox2 are expressed different cells.**

a. Widefield images of Nos2, Cox2, α-SMA, and DAPI multiplexed immunofluorescence staining surrounding a blood vessel in a 4T1 tumor.
b. Widefield images of Nos2, Cox2, and DAPI immunofluorescence staining of a 7-day old 4T1 disk.
c. Nos2 vs. Cox2 intensity for each cell in the 4T1 tumor section shown in Supplemental Fig. 9a (Pearson's correlation coefficient $\rho$=0.2477).
d. Nos2 vs. Cox2 intensity for each cell in the 4T1 disk shown in Supplemental Fig. 9b (Pearson's correlation coefficient $\rho$=0.0969).

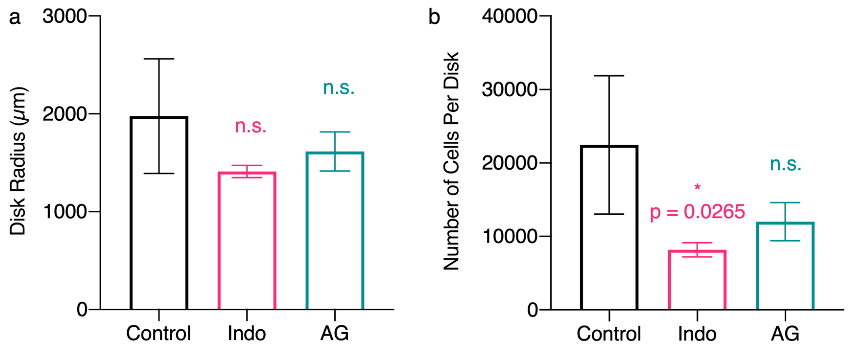


**Supplemental Figure 10. Effect of treatment on 4T1 disk size.** 4T1 disks treated with indomethacin have significantly fewer cells than control 4T1 disks, but there is no significant difference in 4T1 disk radius after Indo or AG treatments.

a. Average radius of control disks and disks treated with indomethacin or aminoguanidine.
b. Average number of cells per disk in control disks and disks treated with indomethacin or aminoguanidine.

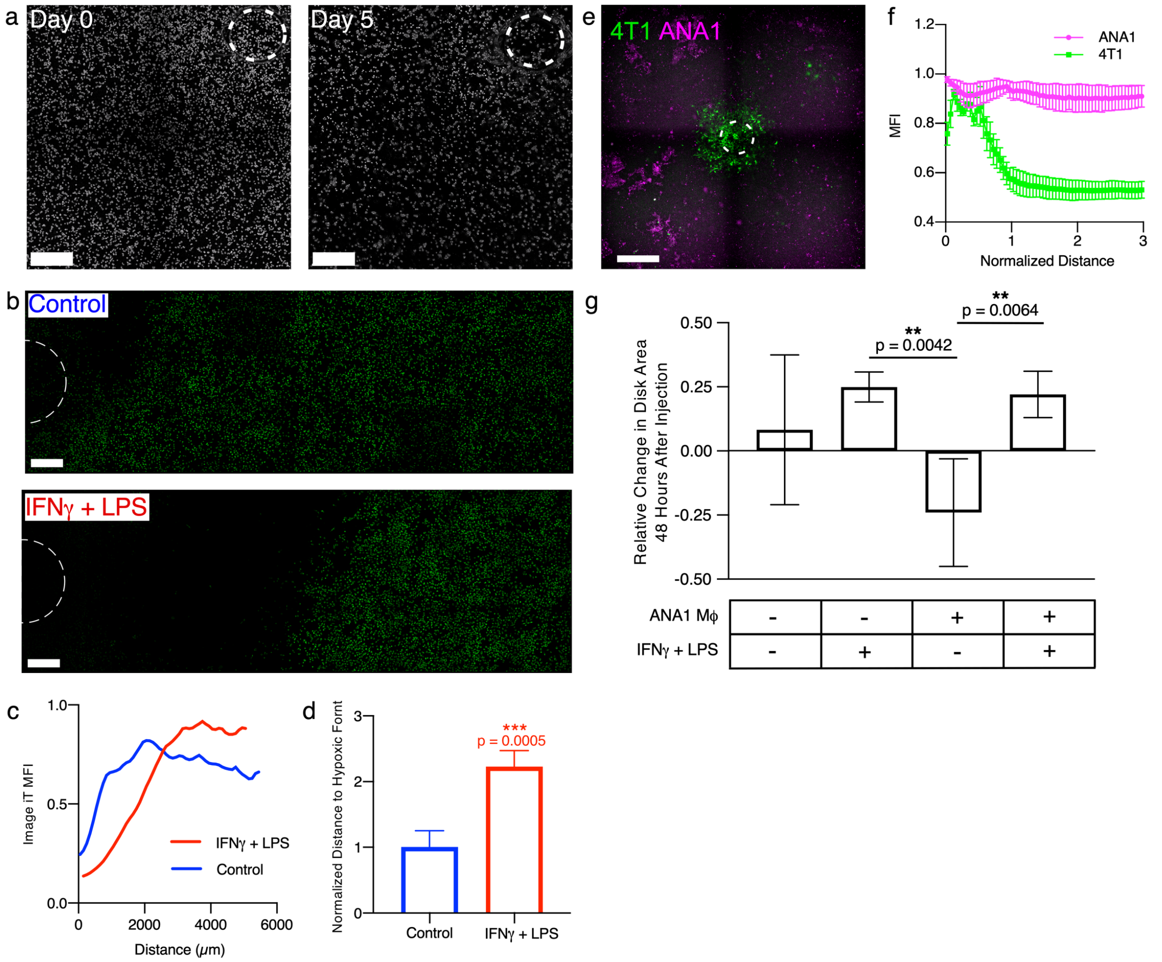


**Supplemental Figure 11. ANA1 macrophages survive in hypoxic regions of REECs and do not form disks, both alone and in co-culture with 4T1 cells. ANA1s stimulated with pro-inflammatory cytokines push back the hypoxic front, decrease oxygen consumption, and infiltrate the hypoxic core of spheroids.**

a. Phase contrast imaging of ANA-1s in chambers upon initial chamber placement and 5 days after chamber placement. There is very little net movement of the cells (scale bar = 500 μm).
b. Widefield images of Image IT Green Hypoxia Reagent (IGHR) fluorescence, in ANA1-REECs 2 hours after chamber placement, with and without 18 hours of cytokine stimulation by IFNγ+LPS prior to chamber placement (scale bar = 500 μm).
c. IGHR fluorescence intensity in ANA1-REECs as a function of distance from the hole (data smoothed using a 5-point window in GraphPad Prism to minimize oscillations resulting from a technical artefact caused by stitching of image tiles.).
d. Quantification of the distance to the hypoxic front (the distance at which IGHR fluorescent reaches 90% of its maximum), normalized to the unstimulated control. ANA1s treated with IFNγ+LPS increase the distance to the hypoxic front 2-fold compared to untreated ANA1s (N = 4).
e. Widefield image of a 1:1 co-culture of GFP expressing 4T1s and ANA1 macrophages labeled with Cell Tracker Red CMTPX, 7 days after chamber placement (scale bar = 1 mm). ANA1s persist in the hypoxic regions of the chamber where 4T1s have died.

f. Quantification of ANA1 and 4T1 distribution 7 days after chamber placement (N = 3).
g. Quantification of the change in GFP-4T1 disk area 48 hours after injecting media with or without ANA1s, and with or without cytokines IFNγ+LPS. Injection of unstimulated ANA1s caused disk size to decrease, because the addition of oxygen consumers to the chamber increased the size of the necrotic zone. In contrast, injection of stimulated ANA1s caused an increase in disk size comparable to fresh media; this indicates that stimulated ANA1s decrease oxygen consumption and thereby shrink the necrotic zone.

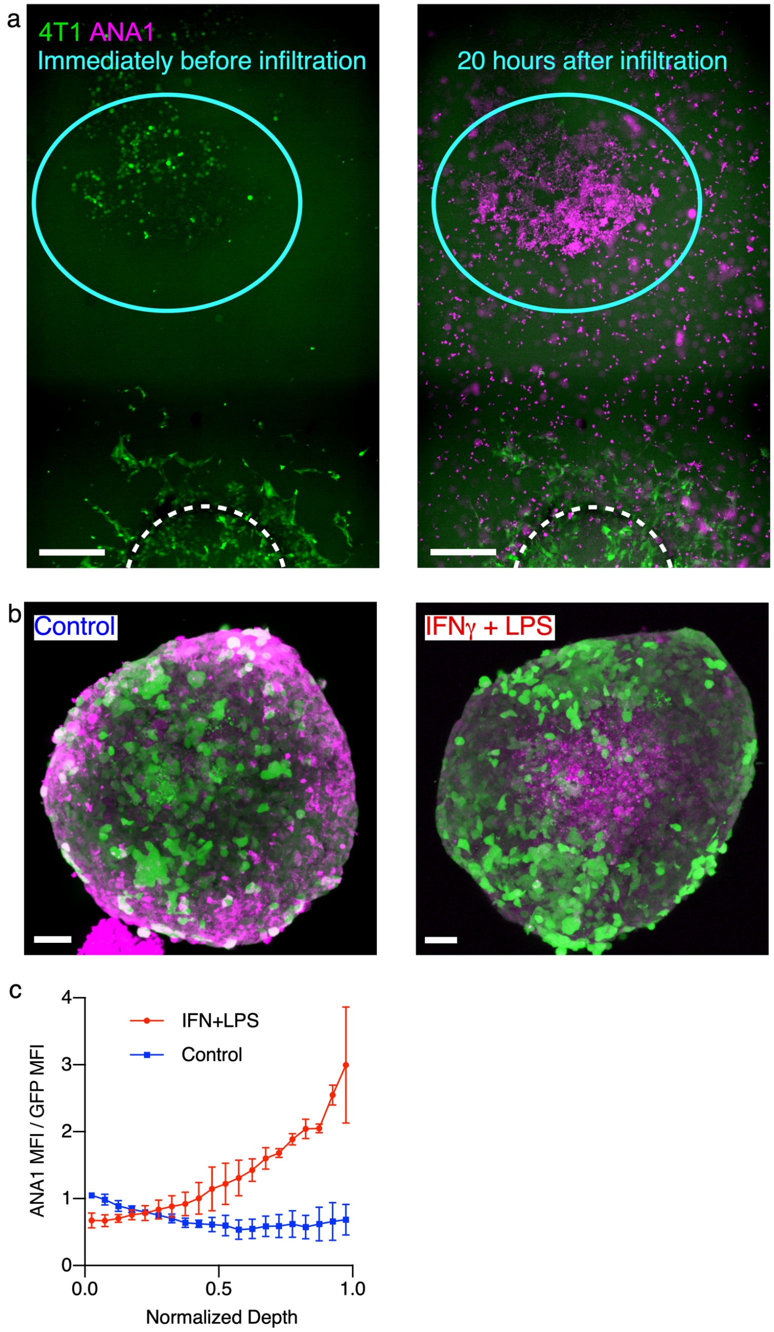


**Supplemental Figure 12. ANA1 macrophages survive in the hypoxic and necrotic of 4T1 disks, and stimulated ANA1 macrophages infiltrate the hypoxic core of spheroids and decrease oxygen consumption.**

a. After 4 days, GFP-4T1 cells form a disk, while some dying GFP-4T1s (outlined with blue dashed circle) remain in the hypoxic region of the chamber, far from the opening (left). ANA1 macrophages were injected into the chamber through the hole. Within 20 hours after macrophage (red) injection, the macrophages are localized to the necrotic zone (right) (scale bar = 700 μm).

b. Confocal microscope images of the equatorial planes of cleared GFP-4T1 spheroids with unstimulated (left) ANA1 macrophages and IFNγ + LPS stimulated macrophages (right) demonstrate stimulated macrophages infiltrate the hypoxic core of 4T1 (green) spheroids (scale bar = 50 µm).
c. Quantification of the ratio of ANA1 mean fluorescent intensity to GFP mean fluorescent intensity as a function of depth from the surface to the center of the spheroids (N = 2 per treatment group).

### Supplemental Protocol

#### REEC construction and use

##### Materials

- Compressed air drill (400x High Speed Engraver, SCM Systems, Menomonee Falls, WI)
- UV Curable epoxy (NOA81, Norland Products, Cranbury, NJ)
- UVO-Cleaner Model 342 (Jelight Company Inc., Irvine, CA)
- 100µm tall 18 mm stainless steel O-rings (PN 90214A123, McMaster-Carr, Robbinsville, NJ)
- 18 mm diameter coverslips, number 1 thickness
- Glass slides
- Scotch tape
- Ethanol
- Forceps (Ted Pella, Redding, CA)
- 500 µm drill bits (SCM Systems)
- 12-well glass bottom plates (PN P12-1.5H-N, CellVis, Mountain View, CA)
- Laser cut mylar clamps
- Laser cut mylar 12-well plate rims

##### Warnings

Make sure to wear appropriate personal protective equipment, including eye and ear protection during glass drilling.

#### Making chambers and plates

##### Modify the multiwell plate

1. Apply UV curable epoxy carefully around each well on a 12-well, glass-bottom plate
2. Apply rims so that each well has even overhang of mylar
3. Cure in UV box for five minutes
4. Use immediately for cell culture or store somewhere covered and protected, sterilize in UV box for at least three minutes before use

##### Assemble the REEC

1. Tape a 18 mm coverslip to a clean glass slide to prevent the coverslip from moving during the drilling. Tape applied to opposite edges of the coverslip, or to 3 equally space positions around the edge is usually sufficient.
2. Drill a hole through the center of the coverslip. Hold the drill normal to the glass surface and press lightly with the drill on the glass -- do not push hard.
3. Once the drill has gone through the coverslip, remove the coverslip from the slide. Clean debris from the coverslip with kimwipes and ethanol. It is important to ensure there are no pieces of glass or tape residue on the coverslip.
4. Apply UV curable epoxy carefully to a stainless-steel O-ring, place glass coverslip using forceps on the O-ring. Avoid applying too much epoxy.
5. Cure in UVO-Cleaner for five minutes, glass side up.
6. Apply glue around clamp

7. Place glued coverslip and O-ring on the clamp so that the glass is in between the mylar clamp and the stainless-steel O-ring
8. Cure in UVO-Cleaner for five minutes, mylar side up.
9. Flip over with forceps
10. Place in UVO-Cleaner for three minutes.
11. At this point, the chambers are sterile. Use immediately or store in covered location and sterilize in UVO-Cleaner for at least 3 minutes, each side, before use.

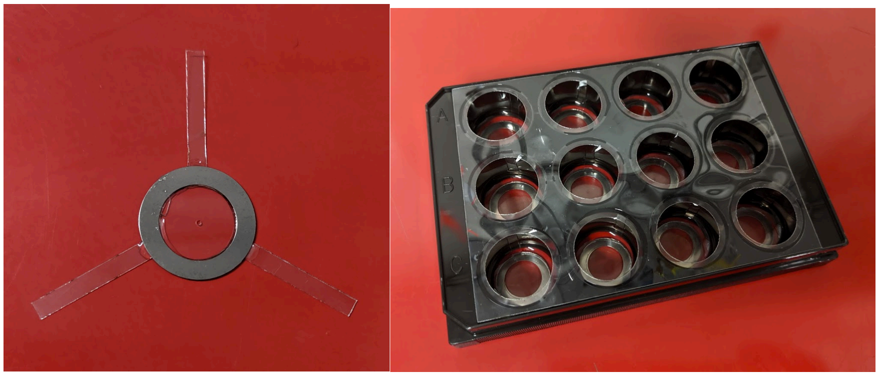

**REEC assembly. Left:** Fully assembled REEC, including coverslip with through hole, mylar clamps and O-ring. **Right:** Fully assembled REECs in a well of a 12-well plate with PBS.

**Cell Culture for REECs**

1. Plate 1 mL of cells per well with desired density of cells in sterile, rim-glued, 12-well, glass-bottom plate. For 4T1s, use a density of 400,000 cells/mL.
2. Allow cells to adhere for an appropriate time. This will vary by cell type. For 4T1s, allow to adhere overnight.
3. To place chamber, use forceps to gently push sterilized chamber down onto cells. Avoid scratching surface and removing cells or allowing air bubbles to form in the chamber.
